# Overcoming nucleotide bias in the nonenzymatic copying of RNA templates

**DOI:** 10.1101/2024.09.03.610991

**Authors:** Daniel Duzdevich, Christopher E. Carr, Benjamin Colville, Harry R.M. Aitken, Jack W. Szostak

## Abstract

The RNA World hypothesis posits that RNA was the molecule of both heredity and function during the emergence of life. This hypothesis implies that RNA templates can be copied, and ultimately replicated, without the catalytic aid of evolved enzymes. A major problem with nonenzymatic templated polymerization has been the very poor copying of sequences containing rA and rU. Here we overcome that problem by using a prebiotically plausible mixture of RNA mononucleotides and random-sequence oligonucleotides, all activated by methyl isocyanide chemistry, that direct the uniform copying of arbitrary-sequence templates, including those harboring rA and rU. We further show that the use of this mixture in copying reactions suppresses copying errors while also generating a more uniform distribution of mismatches than observed for simpler systems. We find that oligonucleotide competition for template binding sites, oligonucleotide ligation, and the template binding properties of reactant intermediates work together to reduce product sequence bias and errors. Finally, we show that iterative cycling of the activation chemistry and templated polymerization improves the yield of random-sequence products. These results for random-sequence template copying are a significant advance in the pursuit of nonenzymatic RNA replication.

## INTRODUCTION

The RNA World hypothesis for the emergence of cellular life implies that RNA molecules can replicate in the absence of evolved enzymes (1). Primordial replication would have been necessary to propagate the information encoded by RNA for the emergence of a ribozyme-based form of primitive life (2). The nature of the protogenetic information and the mechanism of replication remain unknown. Any such replicative process would have required template-directed nonenzymatic polymerization to copy the information stored in a given RNA template. Laboratory models of templated copying use chemically-activated nucleotides and short defined-sequence templates (3,4). However, an ideal nonenzymatic replication mechanism should be able to copy arbitrary sequences, as observed for evolved DNA genome replication. Previous work with all four canonical ribonucleotides as reactants (5) showed that copying random-sequence templates is inefficient compared with copying defined templates. Copying random-sequence templates is heavily biased to rG and especially rC products, and products are also generally error-prone (0.11 frequency of incorrect incorporations). That work and the results presented here both use a nucleotide activation chemistry that results in the formation of reactive bridged dinucleotides (see below), and a higher proportion of bridged dinucleotides to activated mononucleotides is known to correlate with a lower error frequency. The inclusion of methyl isocyanide (MeNC) activation chemistry gives a higher-than-equilibrium proportion of these bridged dinucleotides, and therefore reduced the error frequency to 0.073, but did not improve the product sequence biases. Despite the preference for rG/C-rich templates, all 64 possible sequence products of at least three bases in length were represented, suggesting that although some sequences are heavily favored and others disfavored, all can in principle be accessed by copying.

In this study we tested whether the inclusion of random-sequence 5′-phosphorylated oligoribonucleotides of different lengths in the template copying reaction mixture could reduce the sequence bias of products. Such a mix of random-sequence oligos would be a plausible part of a natural origin of life scenario. We anticipated that additional oligos may reduce bias primarily by participating in the formation of reactive mononucleotide-bridged-oligonucleotide species that should bind a range of template sequences with higher stability than bridged dinucleotides alone. In fact, many reactant species and intermediates are expected to participate in templated polymerization when oligonucleotides are included (Figure 1A-H). We also incorporated *in situ* MeNC activation chemistry. We chose to work with the well-established and prebiotically plausible 2-aminoimidazole (2AI) activating group (Figure 1A) (3,6,7). 2AI-activated mononucleotides form a reactive 5′-5′ phospho-imidazolium-phospho bridged dinucleotide (N*N) in solution at a pH approximately equal to the pK_a_ of the imidazole (pH ∼8 for 2AI-p) (8). Bridged dinucleotides can bind a primer-template complex by Watson-Crick base-pairing (9). The 3′-OH of the primer then attacks the proximal phosphate, displacing an activated mononucleotide as the leaving group. The primer is consequently extended by +1, even though the bridged dinucleotide intermediate consists of two nucleotides. Direct primer extension *via* the activated mononucleotide is also possible (Figure 1B), but is slow with imidazole-based activation, and error-prone compared to the bridged dinucleotide mechanism (5). A downstream oligonucleotide can sandwich any reactive species against the primer and improve primer extension by preorganizing the local helical structure of the RNA (Figure 1C) (10–13). An activated mononucleotide can also form a 5′-5′ bridged species with an activated oligonucleotide (Figure 1D) (11,14). We have previously termed these “helper oligos.” Such mononucleotide-bridged-oligonucleotides are analogous to bridged dinucleotides, except that the downstream component is an oligo instead of a mononucleotide, and therefore better able to bind the template due to lower *k*_off_ rates. Activated oligonucleotides can ligate directly to the primer (Figure 1E). This is known to be a slower reaction than that with bridged dinucleotides or mononucleotide-bridged-oligonucleotides (11,15). Oligonucleotides can also ligate *via* a sterically displaced bridged mononucleotide (Figure 1F) (14). Here, a 5′-5′ bridged mononucleotide-oligonucleotide binds with the oligonucleotide flush against the primer, and the leaving group is an activated mononucleotide. This reaction is faster than direct ligation of an oligonucleotide, i.e. with just the imidazole as the leaving group, as in Figure 1E. Finally, many different species will compete for binding sites in the context of a heterogeneous reaction, and this dynamic may affect the distribution of product sequences (Figure 1G).

**Figure 1.**
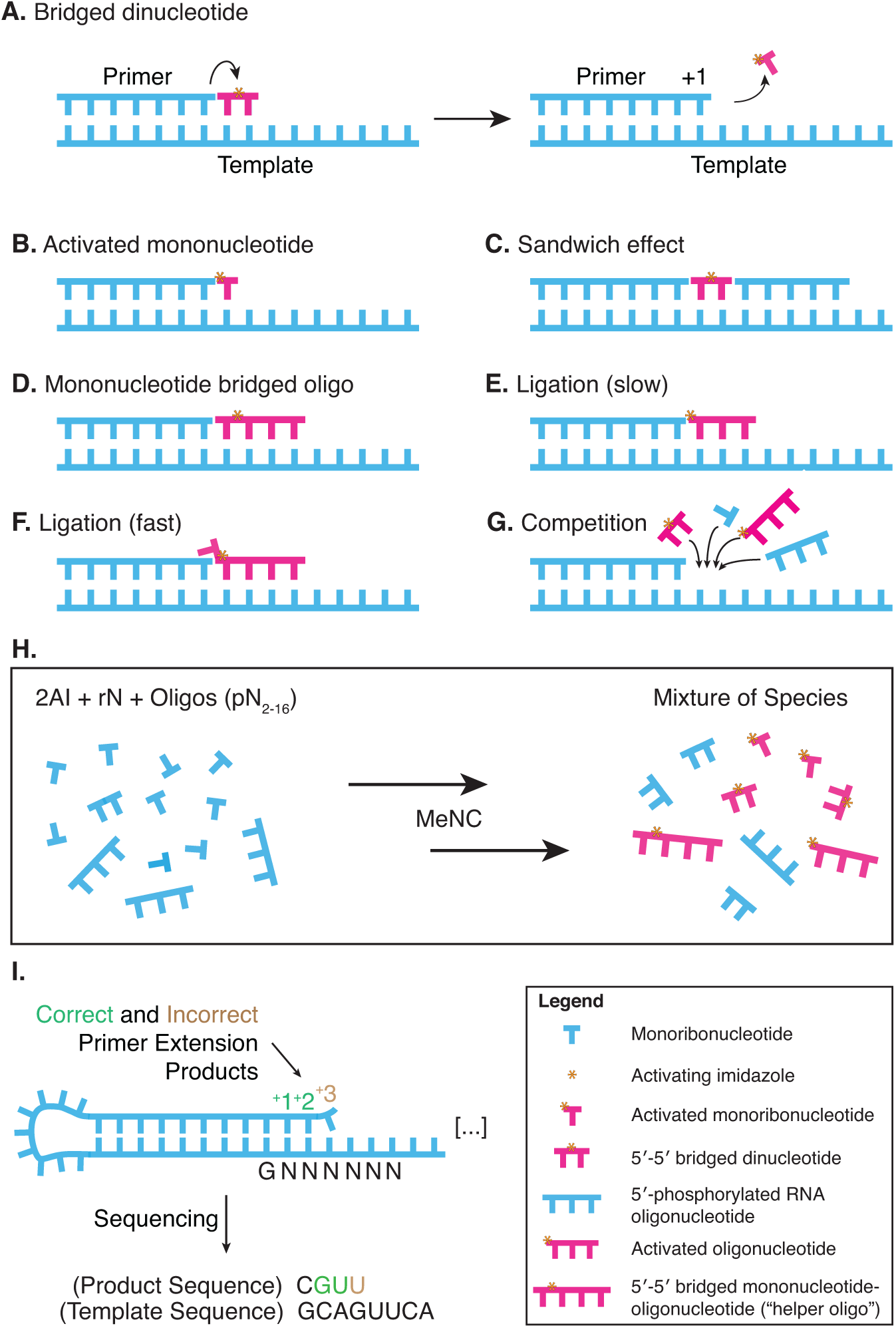
Templated nonenzymatic polymerization, activation chemistry, and sequencing of products. **A.** Mechanism of templated polymerization by imidazole-activated 5′-5′ bridged dinucleotide that can bind a primer-template junction. **B.** Direct extension *via* the activated mononucleotide. **C.** A downstream oligonucleotide can sandwich any reactive species against the primer. **D.** A mononucleotide can form a 5′-5′ bridged species with an oligonucleotide. **E.** Direct ligation of an activated oligonucleotide, analogous to mechanism (B). **F.** Ligation *via* a sterically displaced 5′ bridged mononucleotide. **G.** Many species compete for binding sites in a heterogeneous reaction. **H.** A mixture of mono- and oligonucleotides can be activated *in situ* by MeNC-mediated chemistry. **I.** (Adapted from 1.) A self-priming hairpin with a random-sequence template captures the products of polymerization. Sequencing analysis then reveals the ensemble of template-specific products.

Previous work has shown that methyl isocyanide (MeNC), in combination with a simple aldehyde such as 2-methylbutyraldehyde (2MBA), can drive *in situ* activation of mononucleotide or oligonucleotide 5′ phosphates with imidazole (16). Nucleotide activation can be achieved with equimolar concentrations of an imidazole by repeated freezing of all components in eutectic ice (17). Cyclical freeze-thaw could have occurred on the early Earth or early Mars (18,19). Under these conditions of heterogeneous ice nucleation, the nucleotides and activating agents are thought to become concentrated in microscopic channels that remain aqueous in the context of solute-excluding ice (20). Subsequent solution-phase activation by the further addition of MeNC and 2MBA or other simple aldehyde promotes the formation of 5′-5′ bridged species (21,22). Here, we use this MeNC-driven eutectic activation followed by solution-phase activation to generate complex mixtures of activated mononucleotides, oligonucleotides, and bridged species (Figure 1H), allowing us to access all the pathways shown in Figure 1A-G in a single reaction.

This study required the characterization of sequences from an ensemble of arbitrary templates and associated products. We used NonEnzymatic RNA Primer Extension sequencing (NERPE-seq) to collect data on millions of individual template-product pairs from each experiment (Figure 1I and Supplementary Figure 1) (23). The sequencing construct is a self-priming RNA hairpin with a random-sequence template (typically 18 bases in this study) that captures the products of primer extension. Sequencing analysis then reveals the ensemble of template-specific products.

By combining a heterogeneous mixture of reactants, prebiotically plausible activation chemistry, and a purpose-developed sequencing technology we were able to determine that random-sequence oligonucleotides in a mixture of lengths from 2-16 promote unbiased copying of arbitrary templates. The effect depends on using both the oligonucleotide mixture and the *in situ* MeNC activation chemistry. We were also able to begin addressing the problem of low product yields by cycling the full reaction multiple times. Taken together, our results reveal a conceptually straightforward solution to the long-standing issue of sequence bias in the nonenzymatic copying of arbitrary RNA template sequences.

## MATERIAL AND METHODS

### General

Reagents, sequencing hairpin syntheses, the nonenzymatic RNA primer extension sequencing (NERPE-Seq) protocol, and the data analysis code package were as previously described (5,23). We summarize the general procedures here in condensed form (adapted from (5)). The Extended Material and Methods in the Supplementary data includes additional details about oligonucleotide synthesis and the preparation of activated nucleotides. Supplementary Table 2 lists all oligonucleotides used in this study.

HEPES buffer was prepared from the free acid (Sigma-Aldrich), adjusted to pH 8 with NaOH, and filtered. Imidazoles (Sigma-Aldrich or Combi-Blocks) were prepared as 1 M stock solutions in water, stored at −20°C, and brought to pH 8 immediately prior to use. Enzymes used for the preparation of RNA for sequencing were from New England BioLabs (NEB). All incubations at a specified temperature were performed in a Bio-Rad T100 thermal cycler.

Nuclear Magnetic Resonance (NMR) ^1^H spectra were acquired on a Bruker Avance III HD 400 MHz Nanobay equipped with a Bruker BBFO probe. All reported chemical shifts (δ) are given in parts per million (ppm) relative to residual solvent peaks, and ^1^H spectra calibrated using the residual solvent peaks relative shift to TMS.

### RNA sample preparation for sequencing (**5**)

Generally, 1 μM sequencing hairpin and 1.2 μM 5’ Handle Block (an oligo that prevents the 5’ end of the hairpin from occluding the template (23)) were annealed in 200 mM buffer by incubation at 95°C for 3 minutes, and cooling to 23°C at 0.2 °C/s. The indicated reactants (typically rN, a mixture of oligonucleotides, 2-aminoimidazole, methyl isocyanide, and 2-methylbutyraldehyde) and 30 mM MgCl_2_ (unless otherwise indicated) were added to initiate the reaction. The total final reaction volume was 30 μl. The mixture was briefly vortexed and incubated at the indicated temperature (typically 23°C) in a thermal cycler with a lid heated to ∼10°C above the incubation temperature to minimize evaporation from and concentration of the reaction mixture. Reactions were quenched with 20 μl 0.5 M EDTA, desalted (G25 NU-CLEAN spin column, IBIScientific, followed by Oligo Clean & Concentrator spin column, Zymo Research), and the NPOM (caged) bases uncaged (24) (385 nm UV exposure for 45 minutes). Samples were PAGE-purified (ZR small-RNA PAGE Recovery Kit, Zymo Research). The RT Handle (template for the reverse transcription primer, Supplementary Table 2) was ligated to the 3’ end of the sequencing hairpin with T4 RNA Ligase 2, truncated KQ. The mixture was treated with Proteinase K, phenol-chloroform extracted, and concentrated with an Oligo Clean & Concentrator spin column.

The RT Primer (Supplementary Table 2) was annealed to the purified sample and the RNA reverse transcribed with ProtoScript II. The mixture was purified with an Oligo Clean & Concentrator spin column. Half the product cDNA was added to a 50 μl Q5 Hot Start High-Fidelity DNA Polymerase PCR reaction with 0.2 μM each of NEBNext SR Primer for Illumina and NEBNext Index Primer for Illumina and run for 6 cycles with a 15 s 62°C extension step. The PCR product was purified by preparative agarose gel electrophoresis (Quantum Prep Freeze ‘N Squeeze spin column, Bio-Rad) and magnetic beads (Agencourt AMPure XP, 1.7:1 volume ratio of magnetic bead suspension to sample volume). Samples were validated and concentrations measured by TapeStation (D1000, Agilent). Paired-end sequencing by MiSeq (Illumina, MiSeq Reagent Kit v3, 150-cycle) produced ∼20 million reads with ∼95% passing the instrument’s quality filter. Previous work has evaluated the effects of 2ʹ-5ʹ linkages on the sequencing protocol and found that they do not measurably alter the reported sequencing data (23,25,26). The experimentally established error associated with each reported base identity is 0.062 ± 0.13% (23). Supplementary Table 3 lists specific conditions for each sequencing experiment. The order in which components are listed corresponds to the order in which they were added.

### Methyl isocyanide synthesis and concentration calculation

Methyl isocyanide was synthesized using a modification of the procedure first reported by Schuster and Casanova (27).

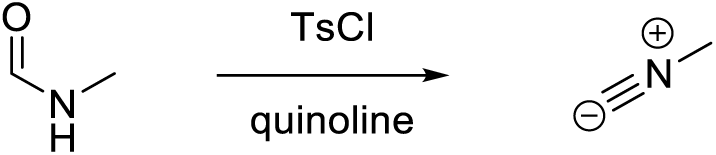

A solution of *p*-toluenesulfonyl chloride (57.2 g, 300 mmol) in freshly distilled quinoline (100 ml) was heated to 75°C in an oil bath, and the system evacuated to a pressure of 15 mm Hg *via* a connected cold finger receiver cooled in a bath of liquid nitrogen. The mixture was stirred vigorously and *N*-methylformamide (11.8 ml, 200 mmol) was added dropwise, to maintain a smooth distillation rate, over 15 minutes. When addition was complete, heating was stopped and the vacuum was released. The product methyl isocyanide (MeNC) was directly collected from the receiver. MeNC aliquots were stored in pre-cleaned 4 ml glass vials with PTFE-backed caps, sealed with electrical tape, and placed in a secondary container at −20°C.

The purity and concentration of methyl isocyanide were measured by ^1^H NMR analysis using trimethyl phosphate as an internal standard. A solution of trimethyl phosphate (0.1 mmol) was prepared in >99% D_2_O (600 μl) (0.165 mM). To this methyl isocyanide (10 μl) was added rapidly, mixed thoroughly, and transferred to an NMR tube. Setting the integration of the methyl peaks of trimethyl phosphate (9 protons) to 3.0, the concentration of methyl isocyanide (3 protons) can be derived as the relative integration given the molar ratio of trimethyl phosphate to methyl isocyanide. Therefore, the amount of methyl isocyanide in a 10 μl aliquot is equal to the integral x 0.1 mmol, and the concentration in mM is the integral x 0.1 mmol divided by 10^-5^ l (13.4 M in the preparation shown in **Supplementary Figure 2**).

### Methyl isocyanide activation of RNA

To avoid pipetting exceedingly small volumes, 1:10 aqueous diluted MeNC working stocks were prepared fresh immediately prior to use. Pipettes and pipette tips for handling the concentrated MeNC stock were cooled to 4°C. 2-methylbutyraldehyde (2MBA, Tokyo Chemical Industry Co.) is poorly soluble in water and was added in one aliquot to the desired final experimental concentration directly from the 9.33 M undiluted stock. Supplementary Table 3 details MeNC activation conditions for each experiment. All experiments with MeNC were performed inside a chemical hood. Direct solution-phase activation was typically with nucleotide reactants, 260 mM 2MBA, 200 mM MeNC, and 200 mM of the indicated imidazole. Eutectic activation followed by solution-phase bridge-forming activation was typically done with nucleotide reactants, 260 mM 2MBA, 50 mM MeNC, and 20.5 mM of the indicated imidazole were added together, flash-frozen in liquid nitrogen, and incubated at - 17°C for 24 hours. The sample was thawed in an aluminum block at room temperature (∼1-2 minutes), and with the addition of the 50 mM MeNC the cycle was repeated for a total of four MeNC additions. After the last frozen incubation and after a cumulative 200 mM MeNC had been added, the solution was incubated at 23°C for 24 hours, followed by the addition of 260 mM 2MBA and 50 mM MeNC. 50 mM MeNC was added at 24 hour intervals until a cumulative 200 mM MeNC had been added, followed by a final 24 hour incubation. Samples were briefly and gently vortexed and spun down at each step in which reagents were added. Each eutectic + solution-phase bridge-forming activation reaction lasts a total of 10 days. The final mixture was quenched in 20 μl 0.5 M EDTA and prepared for sequencing as above. For experiments in which the eutectic activation followed by solution-phase bridge-forming activation was cycled multiple times, the solution was quenched and desalted, and the desalted eluate treated as containing only the sequencing hairpin and associated 5’ Handle Block. All other reactants and reagents were assumed to have been removed by the desalting columns. The rest of the protocol was repeated as for a new experiment, except the annealing step was omitted.

### Sequencing data analysis (**5**)

Sequencing data analysis was performed with the NERPE-Seq custom code package written in MATLAB (MathWorks) and described in (23). Briefly, the code filters raw data using read quality scores, and by checking that reads agree with defined-sequence regions of the hairpin construct and that forward and reverse reads (which overlap) agree with each other. Template and product sequence pairs are extracted and characterized. The template sequence of the construct does not contain precisely equal fractions of rA, rC, rG, and rU (23), so the template from a control experiment in which no activated nucleotides were added was used to generate normalization factors. All presented data are normalized— showing results as they would appear if the template were perfectly random with equal base ratios. The complete descriptive annotation of this code package is documented in a previous publication (23), and the code is publicly available and internally annotated on GitHub (see link at Data Availability).

## RESULTS

### Random-sequence oligonucleotides reduce sequence bias in primer extension products

#### Complementary products are heavily biased to rG- and rC in reactions with all four activated canonical ribonucleotides

We began by reproducing our prior results for template-directed primer extension with 2AI-activated canonical ribonucleotides (2AIrN). These results serve as a standard of comparison for subsequent experiments. We incubated the sequencing hairpin with 20 mM 2AIrN (5 mM each of 2AIrA/U/G/C), 200 mM MeNC, and 260 mM 2MBA for 24 hours at 23°C (Figure 2A-C and Supplementary Figure 3A-D). Under these conditions, with pre-activated nucleotides, MeNC promotes the formation of bridged dinucleotides to a higher peak concentration (∼40%) than they would reach spontaneously without the added activation chemistry (∼18%) (21). Primer extension carried out in this way results in a strong rG/C bias among fully complementary products (*i.e.*: primer extension products with no errors) (Figure 2A-B). The sequence space of primer extension products can be represented as a frequency at a given position where the proportions of each base sum to 1. The frequencies of rA and especially rU in the primer extension products are very low: the average frequency of rA is 0.068 and the average frequency of rU is 0.028 for products across positions +1 through +4. To assess sequence biases more quantitatively, we plotted the frequencies of trimer products (products of at least 3 nucleotides in length) from among fully complementary products (Figure 2C and Supplementary Figure 3D). The highest frequency trimers are NCC, NGC, and NGG in each group as defined by the first product base. The median trimer frequency is 0.002, whereas the ideal frequency for each trimer in a uniform distribution would be 1/64 = 0.016. This strong sequence bias would have posed a problem for copying diverse RNA templates during the emergence of life on the early Earth.

**Figure 2.**
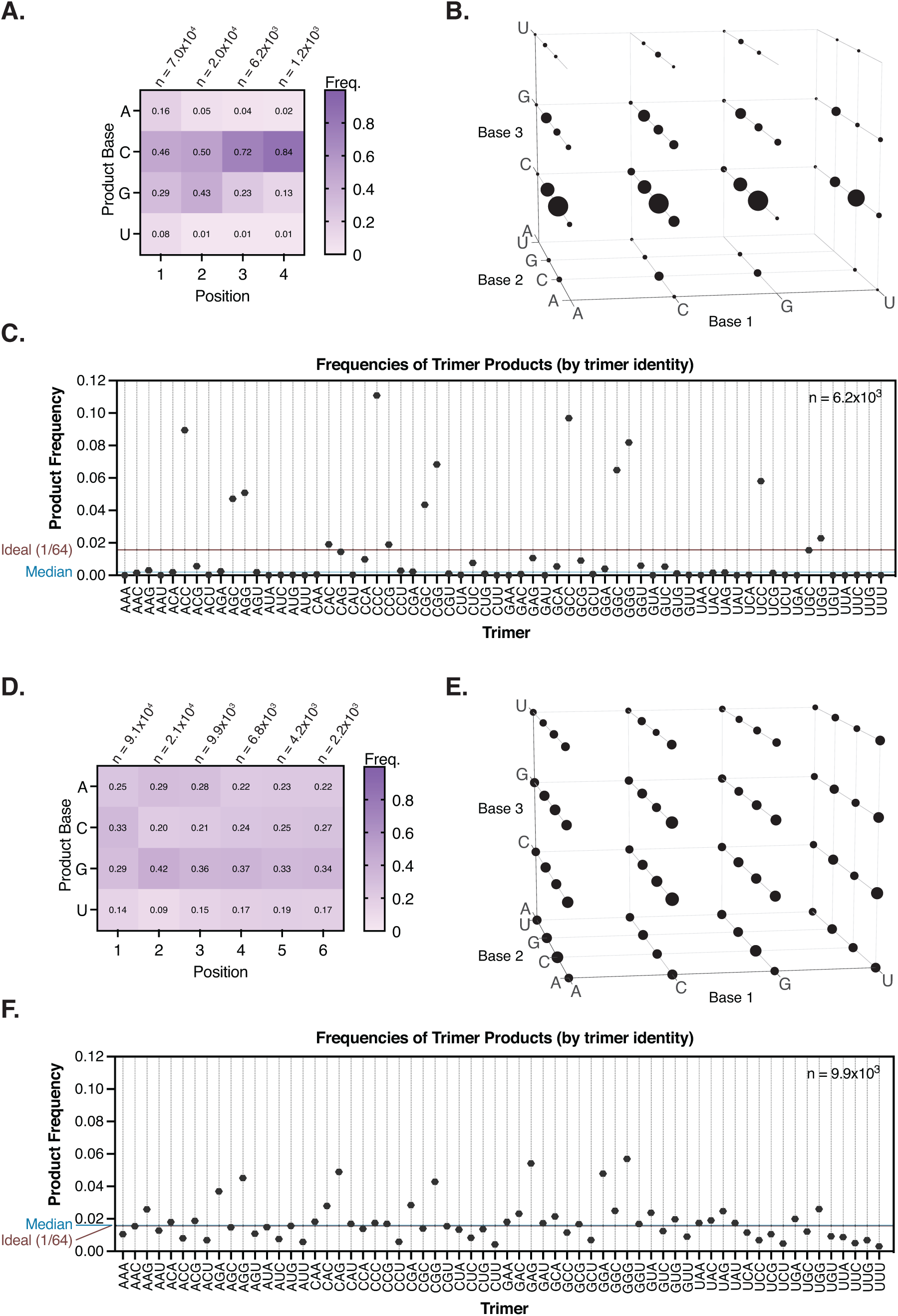
Primer extension with MeNC activation chemistry and a mixture of oligonucleotides reduces the sequence bias of products. **A-C.** Complementary product sequence features of primer extension with 20 mM pre-activated 2AIrN and MeNC activation chemistry to drive bridged dinucleotide formation. **A.** The position-normalized frequency distribution of products is biased to rG and rC. **B.** A visual representation of all products at least three nucleotides long. The volume of each sphere represents the frequency of the indicated product trimer sequence. **C.** Quantification of product trimer frequencies, ordered by trimer identity. The median frequency is 0.002. The ideal frequency for each trimer is 1/64 = 0.016. **D-E.** Complementary product sequence features of primer extension with an *in situ* activated mixture of 20 mM rN + 1 mM oligo mix (pN_2-16_). **D.** The position-normalized frequency distribution of products and **E.** distribution of trimer products are un-biased. **F.** The quantification of trimer frequencies shows a uniform distribution. The median frequency is 0.016. The ideal frequency for each trimer is 1/64 = 0.016.

#### Complementary primer extension products are unbiased in the presence of MeNC activation and a mixture of oligonucleotides

Realistic templated polymerization in a prebiotically plausible setting should include activation chemistry and a mixture of mononucleotides and oligonucleotides, both activated and unactivated. We therefore determined how the inclusion of MeNC activation as well as random-sequence oligonucleotides affects the distribution of complementary primer extension products. We pre-mixed a 5 mM stock of random-sequence 5’-phosphorylated oligonucleotides of lengths 2-16 (oligo mix). We then exposed the sequencing hairpin, 20.5 mM 2AI, 20 mM initially unactivated rN mononucleotides (5 mM each of rA/U/G/C), and 1 mM oligo mix (final concentrations: 500 μM pN_2_, 250 μM pN_3_, 125 μM pN_4_, 62.5 μM pN_5_, 31.3 μM pN_6_, 15.6 μM pN_7_, 7.8 μM pN_8_, 3.9 μM pN_9_, 2 μM pN_10_, 0.98 μM pN_11_, 0.49 μM pN_12_, 0.24 μM pN_13_, 0.12 μM pN_14_, 0.06 μM pN_15_, and 0.03 μM pN_16_) to four rounds of eutectic MeNC activation to drive the formation of 2AI-activated species, followed by four rounds of 23°C solution-phase MeNC activation to promote the formation of bridged dinucleotides and bridged mononucleotide-oligos (for simplicity, we refer to this experiment as “rN + oligo mix + MeNC”). We chose these conditions because bridged species are more reactive than 2AI-activated species that are not bridged. Furthermore, mononucleotide-bridged-oligos bind the template along a longer length than bridged dinucleotides. This may enable them to overcome small differences in binding energies among different sequences.

The inclusion of the MeNC activation and the oligo mix in the primer extension reaction yields a remarkably uniform distribution of complementary product sequences (Figure 2D-F and Supplementary Figure 3E-H). The nucleotide frequencies of products are much less biased than in the absence of *in situ* activation and the oligo mix (Figure 2D). There is a small preference for rG in the primer extension products (see below). rA and rU are well-represented at all positions: the average frequency of rA is 0.25 and the average frequency of rU is 0.15 for products across positions +1 through +6. (compare Figure 2A to 3D). Consequently, the distribution of product trimer sequences is also more uniform (Figure 2E-F and Supplementary Figure 3H). The quantification of trimer frequencies shows a pattern in which NAG, NGA, and NGG are the highest-frequency trimers in each group as defined by the first base, while NCC, NCU, NUC, and NUU are the lowest. The distribution is much closer to the median than in the absence of *in situ* activation and the oligo mix (compare Supplementary Figure 3D to 3H). The CCC trimer is the most overrepresented product in the primer extension reaction without the oligo mix, but it shifts to a position in the middle of the distribution in the primer extension reaction that includes the oligo mix. Notably, the measured median frequency is 0.016, equivalent to the ideal 1/64 value.

**Figure 3.**
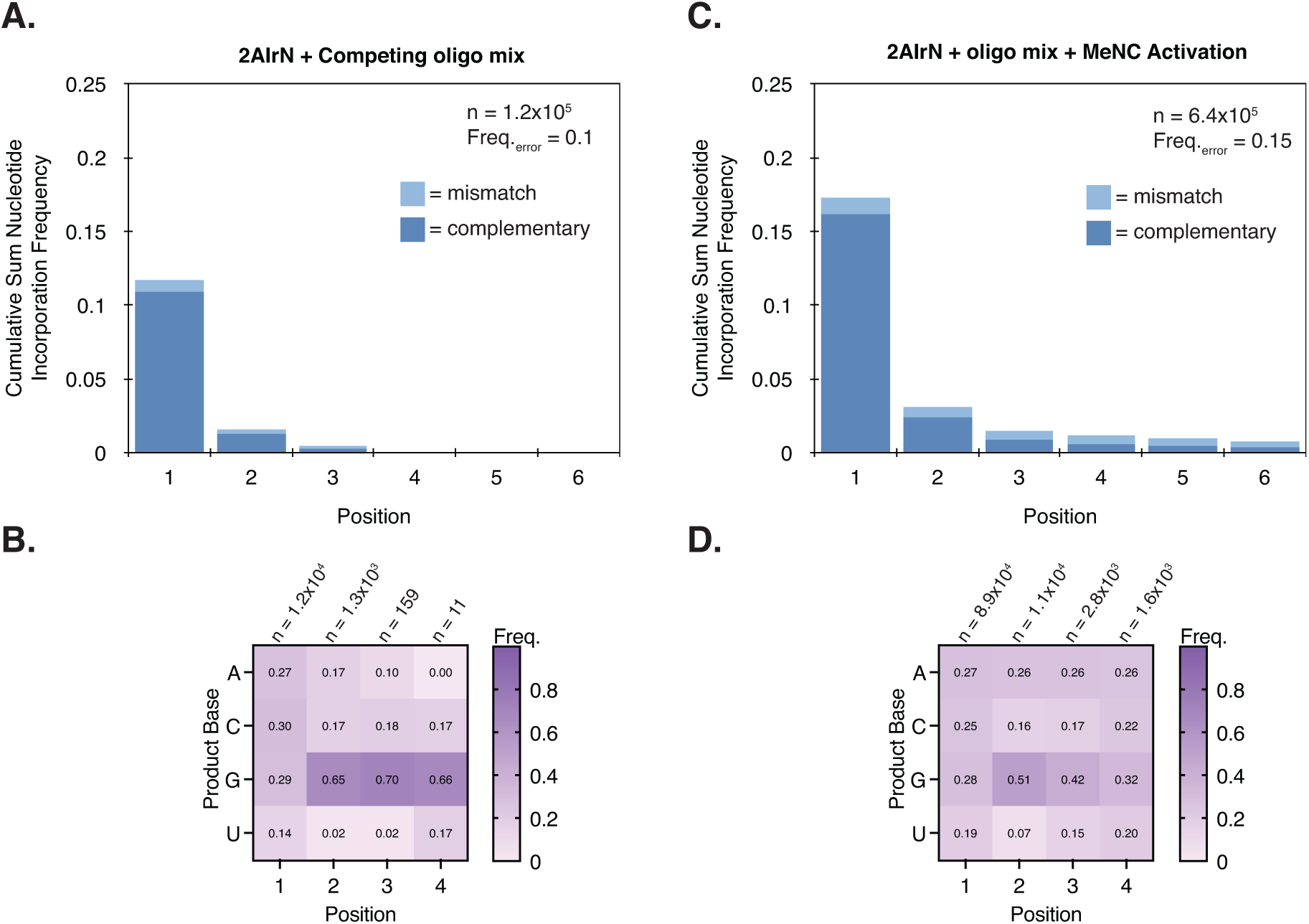
Activated bridged mononucleotide-oligonucleotides are required to maintain yields in the context of competition, and drive an unbiased product distribution. **A-B.** Product sequence features of primer extension with 20 mM 2AIrN + competing unactivated oligo mix. **A.** The cumulative sum frequency distribution of correct and incorrect nucleotide incorporations normalized to each position. The inclusion of the unactivated competing oligo mix suppresses yields and **B.** heavily biases products to rG (see Supplementary Figure 6). **C-D.** Product sequence features of primer extension with 20 mM 2AIrN + initially unactivated oligo mix + MeNC solution-phase activation to drive the formation of activated bridged species. **C.** The cumulative sum frequency distribution of nucleotide incorporations shows higher yields compared to (A). **D.** The distribution of products is more uniform compared to (B).

The inclusion of oligonucleotides in the primer extension reaction clearly affects the sequence of complementary products. To better understand the magnitude of this effect, we quantitatively compared the sequence bias in primer extension experiments without and with *in situ* activation and the oligo mix. In the absence of *in situ* activation and the oligo mix, the lowest-frequency 11 product trimers are not represented at all. The overall distribution of trimers is heavily skewed (Supplementary Figure 3C). The lowest-frequency represented trimer is UAA (0.00013), and the highest is CCC (0.111), an 854-times difference (Supplementary Figure 3D). With *in situ* activation and the oligo mix, the overall distribution of trimers is more normal (Supplementary Figure 3G). The lowest-frequency trimer is UUU (0.003), and the highest is GGG (0.057), a 19-times difference (Supplementary Figure 3H). The yields are similar in both cases, with 20% of all hairpin primers extended without the oligo mix, and 23% of hairpin primers extended with the oligo mix (Supplementary Figure 3A v. 3E). A tail of longer products is evident when *in situ* activation and the oligo mix are included in the primer extension reaction (Supplementary Figure 3B v. 3F). We attribute this to oligonucleotide ligations (see below). Importantly, the inclusion of *in situ* activation and the oligo mix does not compromise fidelity: the frequency of incorrect nucleotide incorporations is 0.063 in both cases.

We considered whether the unbiased distribution of products can be achieved by the inclusion of a set of oligos of one length, rather than the full oligo mix. A primer extension reaction with 20.5 mM 2AI, 20 mM rN, 250 μM pN_3_ trimer oligo (but no oligo mix), and *in situ* activation results in rC-biased complementary products (Supplementary Figure 3I). The distribution of trimer products (products of at least 3 nucleotides in length, as before) remains somewhat biased, though less than with no oligonucleotides at all (Supplementary Figure 3J). The error frequency is 0.15, compared to 0.063 with just mononucleotides and no added oligos, or with the added oligo mix. The use of just the trimer oligo in the primer extension is therefore insufficient to promote unbiased template copying.

#### Products with multiple mismatches are more common in the primer extension reaction with rN + oligo mix + MeNC

Nonenzymatic templated polymerization is generally error-prone because correct nucleotide incorporations rely on correct base pairing without the aid of enzymes that can sense mismatches. We have previously shown that the error frequency of nonenzymatic primer extension depends on the ratio of activated mononucleotides to bridged dinucleotides. Higher ratios of activated mononucleotides correspond to higher error frequencies (5). The experiment described above in which primer extension is performed with 20 mM 2AIrN and solution-phase MeNC activation, but no added oligonucleotides, results in a previously-identified pattern of mismatches (Supplementary Figure 4A). Namely, the A:G mismatch (template:product) dominates position 1, and G:U mismatches are more common at positions ≥2 (5). The distribution of mismatches changes for the rN + oligo mix + MeNC primer extension reaction (Supplementary Figure 4B). The high frequency of A:G mismatches in position 1 is still present, but G:U and U:G mismatches are more highly represented.

Mismatches are typically terminal in reactions with just activated mononucleotides (28). Therefore, in the experiment with 2AIrN but no oligo mix, most polymerization products with a mismatch contain only one mismatch, and that mismatch is the last nucleotide added to the growing primer (Supplementary Figure 4C-D). However, products harboring multiple mismatches are more common in the rN + oligo mix + MeNC experiment (Supplementary Figure 4C-E). Among all products with at least one mismatch, the frequency of products with > 1 mismatch is 0.06 without the oligo mix, and 0.15 with the oligo mix. However, the overall error frequency of incorrect incorporations is the same in both cases (0.063, as reported above), and the majority of products are fully complementary in both cases: 91% without the oligo mix, and 92% with the oligo mix. Below, we show that the higher frequency of products with multiple mismatches in the experiment with the oligo mix is consistent with a contribution from oligonucleotide ligations.

### Unactivated mononucleotides and oligonucleotides do not promote unbiased template copying

#### Unactivated competitor RNAs reduce product yields and fidelity

The rN + oligo mix + MeNC experiment contains a mixture of nucleotide components, including unactivated and activated species of different lengths, all of which may contribute to which sequences are copied. We sought to systematically establish the contributions of each of these components to the resulting primer extension product sequences. Experiments with MeNC use either periodic or only initial activation of nucleotides. In both cases, this allows for periods during which the concentrations of unactivated mononucleotides and/or oligonucleotides are high, for example from hydrolysis. Unactivated species can still compete with activated species for template binding sites and thereby potentially inhibit primer extension. To test the effects of unactivated species, we added various mixtures of unactivated oligonucleotides to a standard reaction with 20 mM pre-activated 2AIrN and no activation chemistry. By excluding the MeNC activation chemistry we ensured that the added competitors remained unactivated and could only passively affect the reaction. Unactivated mononucleotide competitors decrease product yields at concentrations above 30 mM, but have no effect on fidelity or the distribution of nucleotides among complementary products (Supplementary Figure 5A-B). The addition of each set of oligos of a given length from the oligo mix—each one added in a separate experiment—does not generally improve yields or fidelity compared to the rN + oligo mix + MeNC reaction (Supplementary Figure 5C). Further, the inclusion of the entire unactivated oligo mix as a competitor against the 20 mM 2AIrN pre-activated mononucleotides significantly reduces yields and increases error frequencies in proportion to concentration. The addition of 2 mM unactivated oligo mix reduces the frequency of extended hairpins from 23% to 8% and increases the error frequency from 0.063 to 0.13, again compared to the rN + oligo mix + MeNC primer extension reaction (Supplementary Figure 5C).

#### Unactivated random-sequence competitor oligonucleotides show a length-dependent effect on product sequence bias

The oligo mix includes oligonucleotides of lengths 2 through 16. Some oligos will be unactivated over the course of a primer extension reaction, even with MeNC activation chemistry, and may compete for template binding sites. We determined the magnitude and length-dependence of this effect by measuring how each set of oligonucleotides of a given length affects the nucleotide frequencies of complementary products if it is added as an unactivated competitor to a reaction with pre-activated 2AIrN mononucleotides, as introduced above. Recall that the distribution of products in the rN + oligo mix + MeNC primer extension reaction is uniform (Figure 2D), whereas in the reaction with just 2AIrN and no MeNC or oligo mix it is rC-biased (Figure 2A). The addition of very short unactivated oligos (pN_2 or 3_) has little effect on the pronounced rC complementary product bias seen with just 20 mM 2AIrN pre-activated mononucleotides. The addition of unactivated pN_4_ results in an intermediate distribution of complementary products with an rC/G bias. The addition of unactivated pN_5,6,7,8, or 9_ results in a pronounced rG bias (Supplementary Figure 6). The addition of unactivated oligos of lengths greater than pN_10_ again results in an intermediate distribution with rC/G bias. Finally, for the addition of unactivated oligos of lengths pN_15_ and greater, the distribution reverts to rC-rich as in the absence of any added unactivated oligos, or with the addition of very short unactivated oligos.

Based on the pattern of complementary product sequences obtained as a function of added competitor oligo length, we infer that competition for template binding sites may explain the different product sequence biases. The added oligonucleotides are unactivated and cannot directly participate in the primer extension reaction either by ligation or the formation of bridged mononucleotide-oligos. The very short oligos interact too weakly with the template to compete with and affect the known binding stability and reactivity of N*C bridged dinucleotides, and therefore have little influence on complementary product sequences. However, longer oligos will bind more strongly, thereby shifting the distribution of available templates so that rC products are less favored. This pattern holds up to a length at which multiple mismatches become statistically very likely for any bound oligo, such that they are no longer effective competitors for template binding.

#### Bridged mononucleotide-oligonucleotides are required for an unbiased product distribution

The results above show that adding sets of unactivated competitor oligonucleotides of one length to primer extension reactions does not yield unbiased complementary products. The addition of the full unactivated oligo mix as a competitor to the reaction with 20 mM 2AIrN pre-activated mononucleotides shows a rG-rich product distribution, as observed above for some of the added competitor oligonucleotides of one length (Supplementary Figure 7 and Figure 3). The unactivated oligo mix also suppresses product yields (Figure 3A). Therefore, adding the full unactivated oligo mix as a passive competitor in a primer extension reaction is also insufficient to yield unbiased products. We considered whether using activation chemistry to introduce mononucleotide-oligonucleotide bridged species would reduce product bias. The addition of solution-phase MeNC activation in a primer extension reaction with 20 mM 2AIrN pre-activated mononucleotides and the oligo mix recovers the yields (Figure 3C). Under these conditions, the solution-phase MeNC drives the formation of bridged dinucleotides and bridged mononucleotide-oligonucleotides. Complementary products are also less sequence biased (Figure 3D) and comparable to the distribution observed in rN + oligo mix + MeNC reaction (compare to Figure 2D). This result shows that bridged mononucleotide-oligos are required to achieve an unbiased product distribution.

### Ligations

#### High-concentration MeNC activation results in high error frequencies due to efficient ligation

MeNC activation drives the formation of bridged species, so we tested whether increasing the concentration of MeNC would improve the extent of template copying. We performed experiments in which initially unactivated mononucleotides and oligonucleotides are activated *in situ* by eutectic freezing with MeNC followed by solution-phase activation, as before (Figure 2D-F). We compared the inclusion of each set of oligonucleotides of one length, or the full oligo mix, but using 400 instead of 200 mM total MeNC each at the eutectic freezing step and the solution-phase step (Supplementary Figure 8). The inclusion of any individual set of oligos of one length results in both lower yields and significantly higher error frequencies than the inclusion of the oligo mix. The lowest error frequency is 0.11 with the inclusion of pN_3_, and the highest is 0.35 with the inclusion of pN_7_. Thus in all cases except with the addition of pN_3_, the error frequencies are elevated when compared with the 0.11 error frequency in the primer extension experiment that includes the oligo mix with 400 mM MeNC, and the 0.063 error frequency in the experiment that includes the oligo mix with 200 mM MeNC (Supplementary Figure 8A). Using 400 mM MeNC with the oligo mix improved yields only negligibly compared to using 200 mM MeNC with the oligo mix, while almost doubling the error frequency. This indicates no overall benefit to the higher concentration of MeNC.

The experiments with mononucleotides + sets of oligonucleotides of one length + 400 mM MeNC show a clear contribution from ligation in the position-dependent histograms of nucleotide incorporation, in addition to a contribution from mononucleotides. For example, in the primer extension experiment that included pN_6_, there is a readily apparent set of products at +6 (Supplementary Figure 8C), and these products are exceptionally error-prone (error frequency with pN_6_ = 0.27). The experiment with 200 mM MeNC and the oligo mix also exhibits longer products from ligations (Supplementary Figure 3F), but these do not have high error frequencies compared to those in the experiment with 400 mM MeNC (Supplementary Figure 8F).

We suggest that the observed difference between activation with 400 mM MeNC and with 200 mM MeNC is due to the formation of a higher concentration of bridged mononucleotide-oligos at 400 mM MeNC. It is possible that bridged mononucleotide-oligos participate in fast ligations by the sterically displaced mononucleotide mechanism shown in Figure 1F. We have shown the important role of bridged mononucleotide-oligos in maintaining favorable yields and unbiased product distribution (Figure 3), but at higher concentrations they evidently participate in error-prone ligations. The extreme heterogeneity of components in these experiments precludes direct concentration measurements of various reactive species, so we can only make relative comparisons. Further work will be required to conclusively determine how different ligation modes affect product outcomes.

#### Oligonucleotide ligations result in a different pattern of mismatches than primer extension with activated mononucleotides

To better understand the results ascribed to ligations above, we carried out experiments in which only ligations can occur, with no contribution to primer extension from mononucleotides. Previous work has shown that pre-activated defined-sequence oligonucleotides can lead to the accumulation of ligated products in reasonable yields (29). For example, a primer extension reaction with 500 μM 2AI-CGG trimer incubated with a GCC template yields 30% primer-ligated product after 24 hours (23). In contrast, direct *in situ* activation of oligonucleotides with MeNC is very inefficient (Supplementary Table 1). We were only able to generate statistically significant ligation yields by using solution-phase activation of oligonucleotides with 200 mM 1-methylimidazole (1MI), 400 mM MeNC, and a 48-hour incubation. We performed ligation-only experiments using these conditions with each set of random-sequence oligonucleotides of one length. Yields are very low in each case (Supplementary Figure 9), while error frequencies are high: between 0.09 for the ligation of pN_4_ and 0.48 for the ligation of pN_10_. In all cases except pN_2_, the highest error frequency is at the 3ʹ-end of ligation products. This suggests a more stringent requirement for correct base-pairing at the primer-template junction than at the distal end of the oligonucleotide in order for an oligo to become ligated.

The length of a ligated oligonucleotide should contribute to the probability that it will harbor a mismatch to a random-sequence template. We measured how the length of the ligated oligonucleotides affects the proportion of fully complementary products versus products with one or more mismatches. The proportion of fully complementary products drops precipitously for ligated oligos of length pN_5_ and longer (Supplementary Figure 10A). The ligation reactions with sets of oligonucleotides longer than pN_10_ did not yield products. We also performed the ligation experiment with the full oligo mix. Ligation of the oligo mix increases the frequency of products with > 1 mismatch compared to the primer extension reaction with rN + oligo mix + MeNC (Supplementary Figure 10B).

Finally, we compared the frequency and distribution of mismatches after primer extension resulting only from ligation to those in experiments that also include mononucleotides. We chose the primer extension experiments in which the oligo mix, pN_3_, or pN_6_ were ligated using 200 mM 1-methylimidazole (1MI), 400 mM MeNC, and a 48-hour incubation as representative examples of ligation-only reactions. These three experiments resulted in similar distributions of mismatches (Supplementary Figure 10C-E). All show high frequencies of G:U and U:G (template:product) mismatches, especially at the primer-ligation junction. The overall distribution of mismatches is more uniform with the oligo mix than with either pN_3_ or pN_6_; it is also more uniform than with 2AIrN pre-activated mononucleotides and no added oligos (Supplementary Figure 4A). In comparison, the distribution of mismatches in the rN + oligo mix + MeNC experiment is intermediate between the distribution with just 2AIrN and the distributions with just ligations (Supplementary Figure 4A-B and Supplementary Figure 10C). This intermediate distribution suggests that both activated mononucleotides and ligations contribute to mismatches in the full heterogeneous reaction.

#### Primer extension by ligation with the oligo mix results in an unbiased distribution of complementary products

We have found that ligations can contribute to the spectrum of products obtained when activated oligonucleotides are included in primer extension reactions together with activated mononucleotides. We therefore considered the nucleotide bias in the complementary product sequences resulting from ligation alone. In the primer extension experiment where the oligo mix is ligated with 200 mM 1MI, 400 mM MeNC, and a 48-hour incubation, the nucleotide frequencies of complementary products are unbiased (Figure 4A). This is in contrast to the primer extension result with only mononucleotides and no contribution from ligations, where complementary products are biased in favor of rG and rC (Figure 2A). However, the ligation of the oligo mix is inefficient (Figure 4B), and the frequency of incorrect incorporations is 0.30 (Figure 4C). The quantification of product trimer frequencies shows high frequencies of NAG, NGG, and to a lesser extent NGA in each group as defined by the first product base (Figure 4D). This is distinct from the high frequencies of NCC, NGC, and NGG for the experiment with just activated mononucleotides and no oligonucleotides. The median frequency of product trimers is 0.016, and the CCC trimer is relatively underrepresented. This is again in contrast to the result with only activated mononucleotides, where the median frequency of product trimers is 0.002 and the CCC trimer is heavily overrepresented (Supplementary Figure 3D).

**Figure 4.**
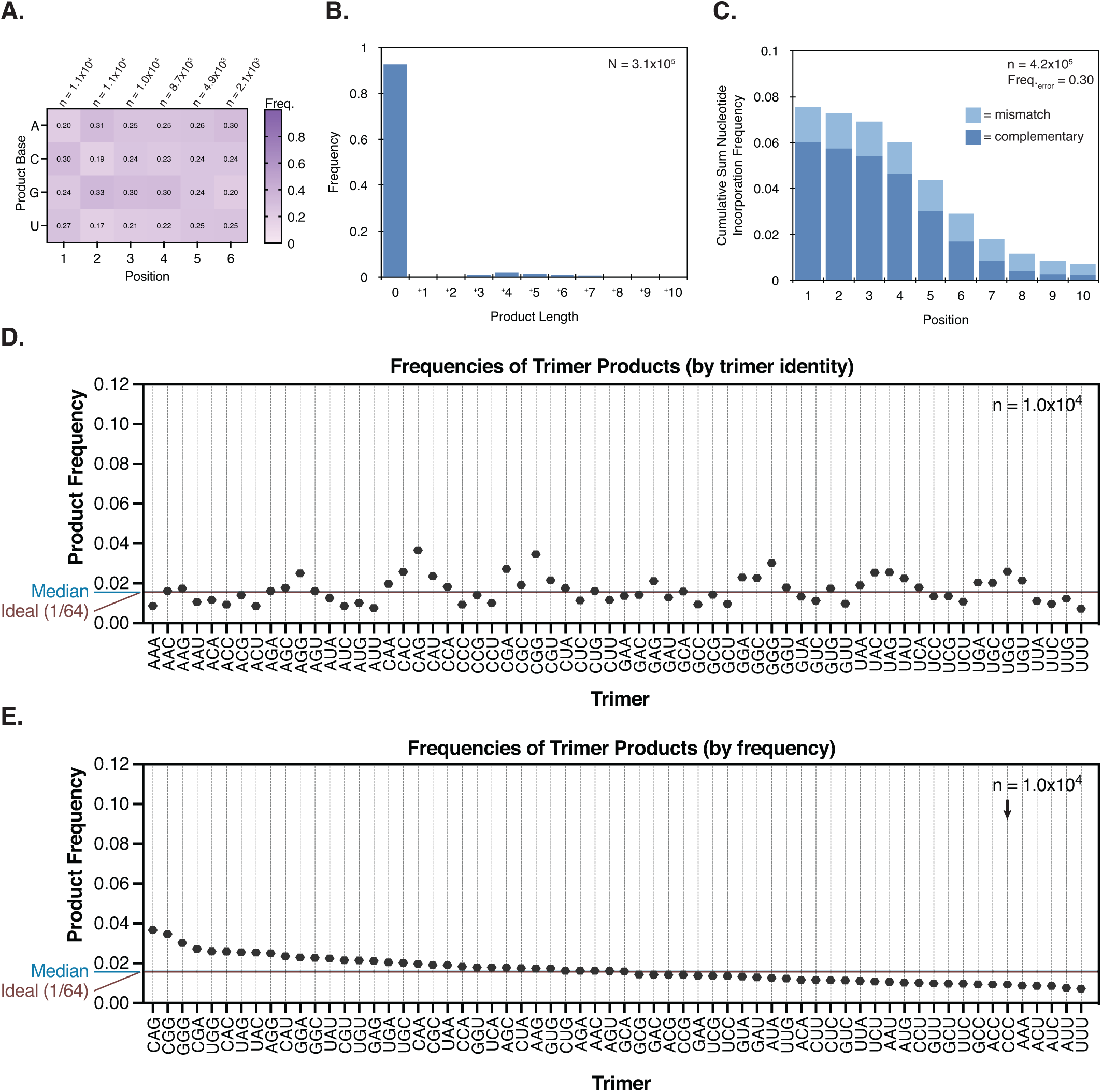
Oligonucleotide ligation-only reactions can be inefficient and error-prone, but exhibit low product sequence bias. **A-E.** Product sequence features of templated ligation with the oligo mix, 200 mM 1-methylimidazole (1MI), and 400 mM MeNC solution-phase activation. **A.** The position-normalized frequency distribution of complementary products is uniform. **B.** Ligations are inefficient under these conditions and **C.** error-prone. **D.** Quantification of product trimer frequencies, ordered by trimer identity. The median frequency is 0.016. The ideal frequency for each trimer is 1/64 = 0.016. **E.** Quantification of complementary product trimer frequencies, ordered by frequency (same data as (D)). Note the relatively flat distribution.

#### Oligonucleotide ligations show length-dependent differences in complementary products

The ligation of the oligo mix described above yields a distribution of complementary products with an unbiased nucleotide composition. We next measured whether this outcome holds with the ligation of sets of oligonucleotides of one length. We again took the ligation of pN_3_ and pN_6_ as representative examples. In contrast to the result obtained with the ligation of the oligo mix, the ligation of pN_3_ yields a relatively biased distribution of products, comparable to that measured for activated mononucleotides alone and no oligonucleotides (Supplementary Figure 11A; compare with Figure 2C and Supplementary Figure 3D). rC- and rG-rich products are overrepresented, and rA- and rU-rich products are underrepresented. Primer extension by CCC, which occurs with the highest frequency in reactions with 2AIrN alone, is here the fourth highest frequency trimer. The median trimer product frequency (products at least three nucleotides in length) is 0.009. In contrast, pN_6_ ligation yields a different nucleotide distribution for positions 1-2-3 that is more uniform and is biased *against* rC- and rG-rich products, with strong representation of rA and rU (Supplementary Figure 11B). The median trimer product frequency is 0.013. The overall trends observed for positions 1-2-3 hold for positions 4-5-6, but here the distribution is even more uniform (Supplementary Figure 11C). The median trimer product frequency for positions 4-5-6 is 0.015.

We interpret the different sequence distributions of complementary products with the ligation of pN_3_ and pN_6_ as a consequence of competition for template binding sites. The product sequence distribution with pN_3_ is similar to that with activated mononucleotides (which form bridged dinucleotides), and no added oligonucleotides. This is consistent with the expectation that trimers behave similarly to bridged dinucleotides, which are effectively dimers from the perspective of template binding. Unlike pN_3_ oligonucleotides, longer oligos such as hexamers are expected to bind a complementary template more strongly, over a longer stretch of nucleotides. Therefore, C/G-rich templates may be more heavily occupied, and any C/G-rich oligo bound at positions 2 or more nucleotides away from the primer can disrupt primer extension by ligation. We find that in the products of primer extension by the ligation of pN_6_, the low representation of rG- and rC-rich product trimers at positions 1-2-3 and 4-5-6 is well explained by the occurrence of ≥2 G, C, or G/C nucleotides in a given product trimer: these occur only 7 times in the top half of the frequency distribution for positions 1-2-3 but 25 times in the lower half (Supplementary Figure 11B). This suggests that multiple templating rG and/or rCs per three-nucleotide window can result in stable enough competitor oligonucleotide binding to disrupt binding of the correct oligonucleotide flush with the primer. Finally, in the experiments described above with pre-activated 2AIrN mononucleotides and unactivated sets of competitor oligonucleotides of one length, we inferred that oligonucleotides of lengths 5-9 preferentially occupy and compete for rG templates relative to rC templates (Supplementary Figure 6). The frequencies of product trimers obtained from the ligation-only experiment with pN_6_ described here enable us to test this inference more quantitatively. Among the top half of the distribution of product trimers at positions 1-2-3 ordered by frequency, the ratio of rC to rG (as the simple fraction of C/G occurrences) is 2.6, whereas in the lower half, it is 0.64. This suggests that rC-rich hexamers are effective at binding rG templates, consistent with our observations with competing unactivated oligonucleotides.

### Cycles of eutectic and solution-phase activation progressively increase yields

Finally, we aimed to begin addressing the problem of low product yields in nonenzymatic templated polymerization reactions, especially in the context of arbitrary sequences. We varied the temperature, timing of MeNC addition, distribution of oligo concentrations in the oligo mix, and oligo lengths used in the oligo mix during the *in situ* eutectic activation and solution-phase activation of reactants anticipating that these variables may affect the hydrolysis of bridged species, or error-prone ligations. However, none of the screened conditions improved on the established 10-day experiment with 24-hour incubations between eutectic freezing steps / MeNC additions, 23℃ solution-phase activation, 20.5 mM 2AI, 20 mM rN, and 1 mM standard oligo mix (Supplementary Figure 12). We therefore iteratively exposed the sequencing hairpin to multiple cycles of *in situ* reactant activation and solution-phase activation under the established conditions. The reaction was quenched and the hairpin isolated between each cycle. The yields improved significantly, with over 69% of all primers extended after six cycles (Figure 5A and Supplementary Figure 13). The frequency of incorrect incorporations is 0.069 for six cycles, compared with 0.063 for one cycle (Figure 5B). The distribution of nucleotides and product trimers among complementary products remain unbiased (Figure 5C-D). The quantification of trimer frequencies for positions 1-2-3, ordered by trimer identity, shows a relatively uniform distribution. NAG, NGA, and NGG trimers remain overrepresented. The median product trimer frequency is 0.014 (Figure 5E and Supplementary Figure 14A). The quantification of trimer frequencies for positions 4-5-6 is also uniform, with a median frequency of 0.014 (Figure 5F and Supplementary Figure 14B). The overall distribution of mismatches is somewhat more uniform than with just one cycle (Supplementary Figure 14C). In summary, our work shows that template copying in the presence of mononucleotides, random sequence oligonucleotides, and activation chemistry overcomes the problem of nucleotide bias, leading to a relatively unbiased distribution of primer extension products. Although the extent of primer extension remains insufficient to copy ribozyme-length templates, further advances in our understanding of prebiotically realistic activation chemistry may lead to a pathway for the continuous nonenzymatic copying of arbitrary templates.

**Figure 5.**
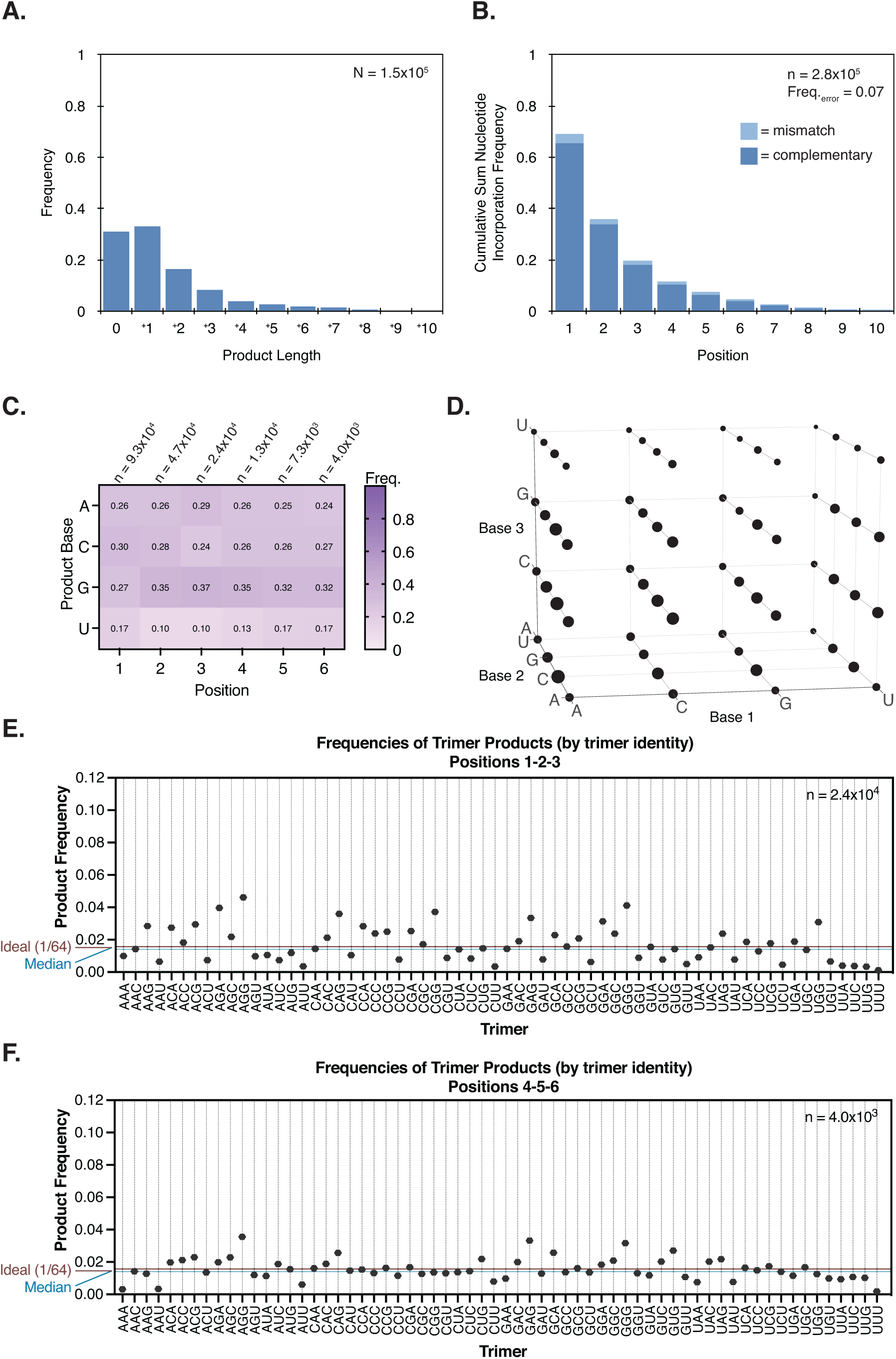
Cycles of eutectic and solution-phase activation increase yields without changing the error frequency or unbiased distribution of products. **A-C.** Product sequence features of primer extension with an *in situ* activated mixture of 20 mM rN + 1 mM oligo mix (pN_2-16_), repeated six times. **A.** The product length distribution shows significantly higher yields after six cycles than just one. **B.** The cumulative sum frequency distribution of correct and incorrect nucleotide incorporations normalized to each position. The frequency of incorrect incorporations is 0.069, compared to 0.063 for one cycle. **C.** The position-normalized frequency distribution of complementary products and **D.** distribution of trimer products remain relatively unbiased, as for one cycle. **E.** The quantification of trimer frequencies for positions 1-2-3, ordered by trimer identity. The median frequency is 0.014. The ideal frequency for each trimer is 1/64 = 0.016. **F.** The quantification of trimer frequencies for positions 4-5-6. The median frequency is 0.014.

## DISCUSSION

We currently lack a self-consistent laboratory model that connects a pool of nucleotide and oligonucleotide building blocks to nonenzymatic templated copying and then to replication. The copying step is central, analogous to the action of DNA polymerases in the context of evolved genome replication. An ideal nonenzymatic copying reaction should be able to copy any sequence, thereby granting emergent proto-evolutionary processes full access to sequence space. Although such an ideal scenario may be an unreasonable expectation in the absence of evolved enzymes, and perhaps not strictly necessary (30), attempts thus far have fallen dramatically short of copying arbitrary sequences without bias. Copying has only been demonstrated for specific defined sequences (3,4,31,32), typically rG/C-rich, and work on arbitrary sequences has shown strong biases (5,33). We do not know how broad a range of possible template sequences must be amenable to efficient copying for a laboratory model of replication and evolution to function. On one end of the spectrum is the specific case, where only one particular sequence can be copied, and on the other is the ideal case, in which any sequence can be copied. The specific case precludes the emergence of evolution because mutations could never become fixed. An intermediate case in which sequences from a restricted region of sequence space can be copied may be an effective scenario. Although the ideal case may seem optimistic, we have shown here that not only can arbitrary sequences be copied by nonenzymatic polymerization in the presence of heterogeneous reactants, but that this heterogeneity is an advantage.

Our results then prompt a follow-up question: is a heterogeneous mixture of reactants plausible? All models of nonenzymatic copying imply at least some initial *un*templated polymerization as a source of templates and primers. It remains unknown whether a successful model system will require only an initial pool of random oligomers, or some background level of periodic or continuous untemplated polymerization. The chemistry of untemplated polymerization may also dictate whether it can happen under conditions that are compatible with templated copying. Further work will be required to determine these variables. In this study we use random-sequence templates and oligonucleotide reactants. We do not know whether random untemplated polymerization can be truly random (34–36), and this will depend on the conditions. Future work will consider how an *in situ*-generated pool of oligonucleotides would affect our reaction. Similarly, we used a distribution of oligonucleotide lengths from pN_2-16_ in decreasing concentrations. This too seems a reasonable assumption for a source of untemplated oligomers, and yielded the best results in our experiments, but ultimately the best test would be the distribution actually generated by some compatible mode of untemplated polymerization.

Reactants such as bridged dinucleotides, oligonucleotides, and bridged mononucleotide-oligos exhibit a broader distribution of template binding energies than just the four individual mononucleotides. This enables otherwise individually unfavored A-U base-pairs to become incorporated as products (4,15,37). The dynamic competition of unactivated oligos for template binding sites exhibits its own bias by occupying rG-rich templates, thereby counteracting the strong preference of the bridged dinucleotide pathway for rC products. Bridged mononucleotide-oligonucleotides seem particularly unbiased when they mediate mononucleotide incorporation. However, they are error-prone when they result in ligation. Finally, ligations of sets of oligonucleotides of one length are somewhat biased, but the ligation of the oligo mix is unbiased. All these pathways work together to smooth out the complementary product distribution relative to the reaction with only activated mononucleotides. Finally, a major pathway we could not address here is the large ensemble of off-template interactions. The components of the complex mixture of oligonucleotides are no doubt interacting among themselves and titrating the relative concentrations of some oligo sequences over others (37). This process is also a natural mechanism to suppress the rC/G bias because rC/G-rich oligos will be likelier to interact with each other off-template.

The inclusion of an *in situ* activation chemistry enabled us to begin addressing the problem of the low yield of primer extension products. More importantly, it opens up new avenues of inquiry about the off-template reactions. The sequencing hairpin construct was developed to understand templated polymerization, but it is not in itself a realistic template. The off-template reactions among the mixture of random-sequence oligonucleotides may be very interesting (38). We anticipate that the phenomena identified here also apply to the reactions that are invisible to us with our current hairpin probe. Future work should develop the sequencing tools required to accurately assess the properties of reactive random-sequence oligonucleotide mixtures.

It is unclear what role a large pool of random templates and reactants may play in the context of growing and dividing protocells. The RNAs outside of the protocells would have access to the bulk volume of reactants, and one can envision the copying of RNA in bulk as a source of oligo feedstocks for the protocells. The relevance of such a process would depend on the permeability of protocell membranes to various-length RNAs; the process of passive endocytosis could allow even long oligonucleotides to slowly enter protocells (39). Large protocells may have significant enough internal volumes to harbor a relatively large and random set of reactants. There will be some tradeoff between using the described mechanisms to copy the specific sequences of the proto-genome in a low-bias way, and having too many longer random oligonucleotides that would scramble the genome. These balances will depend on many additional variables, including the size and nature of the genome (40).

In addition to generating unbiased complementary products, our reaction conditions also result in a more uniform distribution of mismatches relative to the case with only mononucleotide reactants. This would enhance the emergence of an evolutionary process by allowing for a greater variety of mutations. We also determined that cycles of activation and primer extension could be repeated multiple times without compromising the 0.06-0.07 error frequency observed under the most favorable conditions. Although this is orders of magnitude higher than the mutation rate in evolved biological systems, it is comparable to that observed with *in vitro*-evolved RNA polymerase ribozymes. Work with a polymerase ribozyme that uses trimer oligonucleotides as a substrate instead of mononucleotides identified an error frequency of 0.027, while the polymerase ribozyme recently used to replicate another ribozyme for the first time has an error frequency of 0.109 (41,42). Mismatches may also be less likely to propagate in final full-length products because mismatched nucleotides are less likely to prime further polymerization compared to correctly paired nucleotides. This stalling effect may increase the overall effective fidelity of nonenzymatic copying (43).

We found that slow ligation reactions in the context of a mixture of random sequence oligonucleotides can be unbiased and high-fidelity, but ligation reactions that are too fast tend to be error-prone. The ligation rates reported here are relative, but are known to follow the following trend with respect to the leaving group: activated mononucleotide > methylimidazole > 2AI (11,14). Our observed ligation error frequencies correlate with this trend: faster ligations result in more errors. This simple pattern in which fast ligation increases error frequencies is consistent with the faster off-rates of oligonucleotides with mismatches, meaning that faster ligation chemistry is required to capture them (44). The fastest ligation rates are with bridged mononucleotide-oligonucleotides, in which an activated mononucleotide is the leaving group. All conditions tested here that increase the concentration of this species exhibit high error frequencies. However, this same species participates favorably in the reaction when it results in mononucleotide incorporation, driving up yields and reducing bias. This implies that a reactant mixture in which the activation is perfectly efficient, such that all oligonucleotides are activated as mononucleotide-bridged species, may be unfavorable because they could lead to excessive errors through fast ligations. This suggests that a regime in which activation is periodic, which is how these experiments were performed, rather than continuous may be optimal with respect to fidelity. Indeed, the MeNC chemistry is based on a scheme that requires UV light exposure (16), which would vary with day/night cycles.

Additional work will be required to understand how the conditions described here for unbiased template copying may fit into the emergence of a proto-replicative cycle in the context of protocells. Nonetheless, our results reveal a conceptually straightforward and prebiotically plausible pathway to nonenzymatically copying arbitrary-sequence RNA templates with low bias and relatively high fidelity.

## ACKNOWLEDGEMENTS

We thank Aman Agrawal and Aleksandar Radaković for their comments on the manuscript. This article was prepared without the use of artificial intelligence software.

