## Supplementary data for "Overcoming nucleotide bias in the nonenzymatic copying of RNA templates"

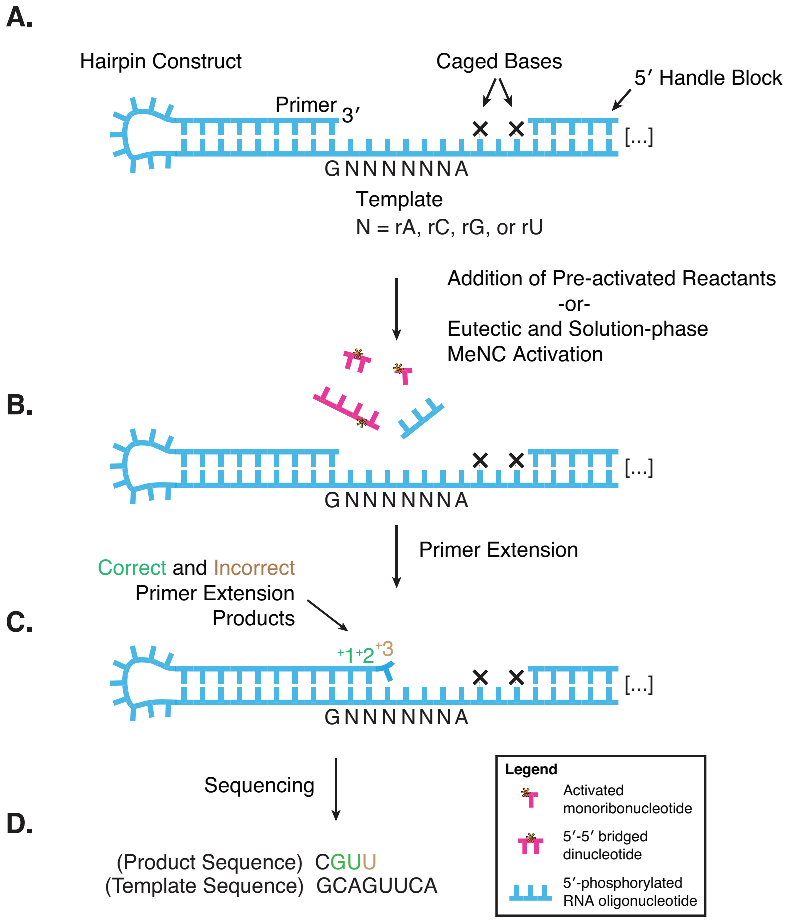

**Supplementary Figure 1. Nonenzymatic RNA Primer Extension-Sequencing (NERPE-Seq).** (Adapted from (1).) **A.** A self-priming RNA hairpin physically links the primer, and any primer extension products, to the random-sequence template (typically 18 nucleotides long in this study). Each sequencing read therefore contains both product and template sequences. The hairpin and the sequence downstream of the random template are defined and used for RNA processing in preparation for sequencing. These defined sequences are also used in the data analysis for quality control and to identify the product and template. Each template terminates in a placeholding "A," which is also useful as a motif for data analysis. The caged bases (2) limit primer extension to a defined position at the end of the template and are uncaged during later processing (3). The 5′ Handle Block oligo base pairs with the 5′ end of the construct to prevent it from interacting with the template (3). **B.** This construct is resilient to methyl isocyanide activation chemistry and can be used directly for all experiments (1). **C.** Each individual hairpin has a specific template sequence that can participate in templated polymerization, yielding correct and / or incorrect nucleotide incorporations at each position. The reaction is quenched, excess reactants are removed, the hairpin is reverse transcribed, and PCR is used to add flanking sequences required for Illumina sequencing. All hairpins are captured for analysis, regardless of the extent of primer extension or absence / presence of errors (3). **D.** Our custom and open-access NERPE-Seq analysis software rigorously filters the data for quality, sorts the sequences from each hairpin into template-product pairs, and performs a range of general analyses on the data (3).

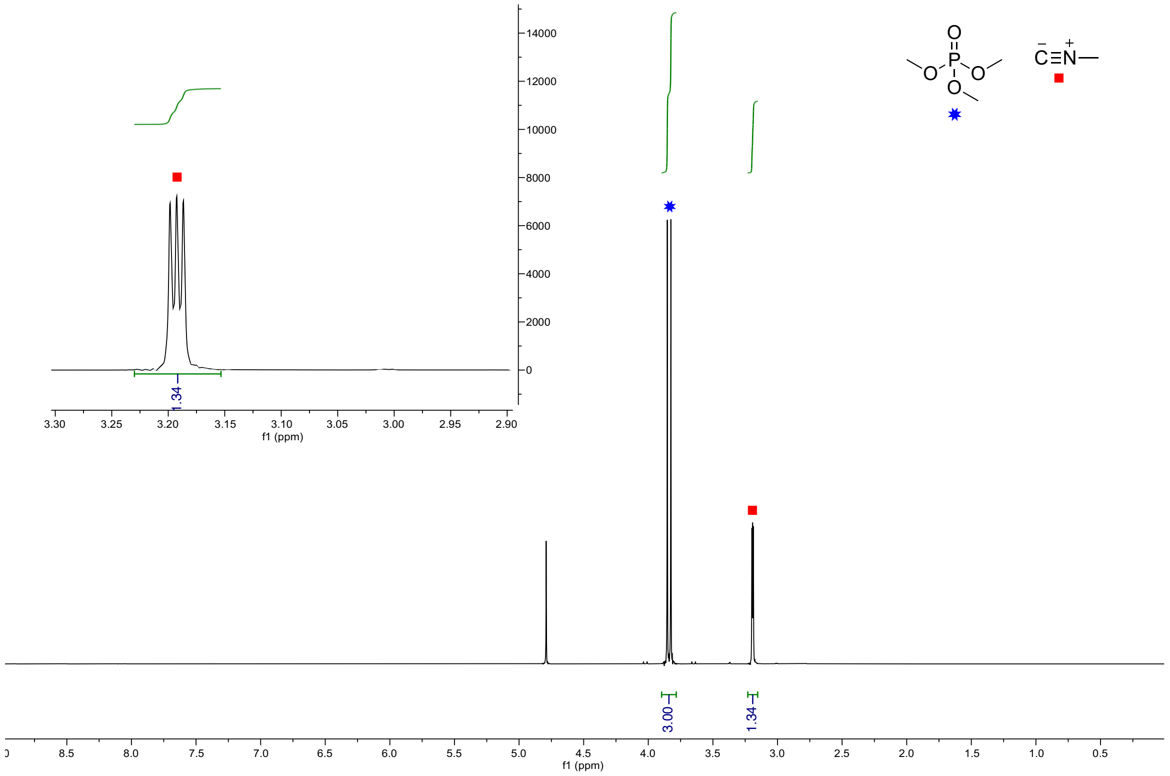

**Supplementary Figure 2. NMR characterization of methyl isocyanide.** The $^{1}H$ NMR (400 MHz, $D_2O$, zg, 0.0-9.0 ppm) spectrum shows methyl isocyanide (■) and trimethylphosphate (✷, internal standard, 3.84 ppm). Inset: expansion of $^{1}H$ NMR (2.90-3.30 ppm).

Methyl Isocyanide

$^{-}C\equiv N^{+}-$

$^{1}H$ NMR (400 MHz, $D_2O$) δ = 3.19 (m, 3H, N-$CH_3$)

Trimethyl Phosphate

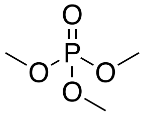

$^{1}H$ NMR (400 MHz, $D_2O$) δ = 3.84 (d, $J$ = 11.1 Hz, 9H, O-$CH_3$)

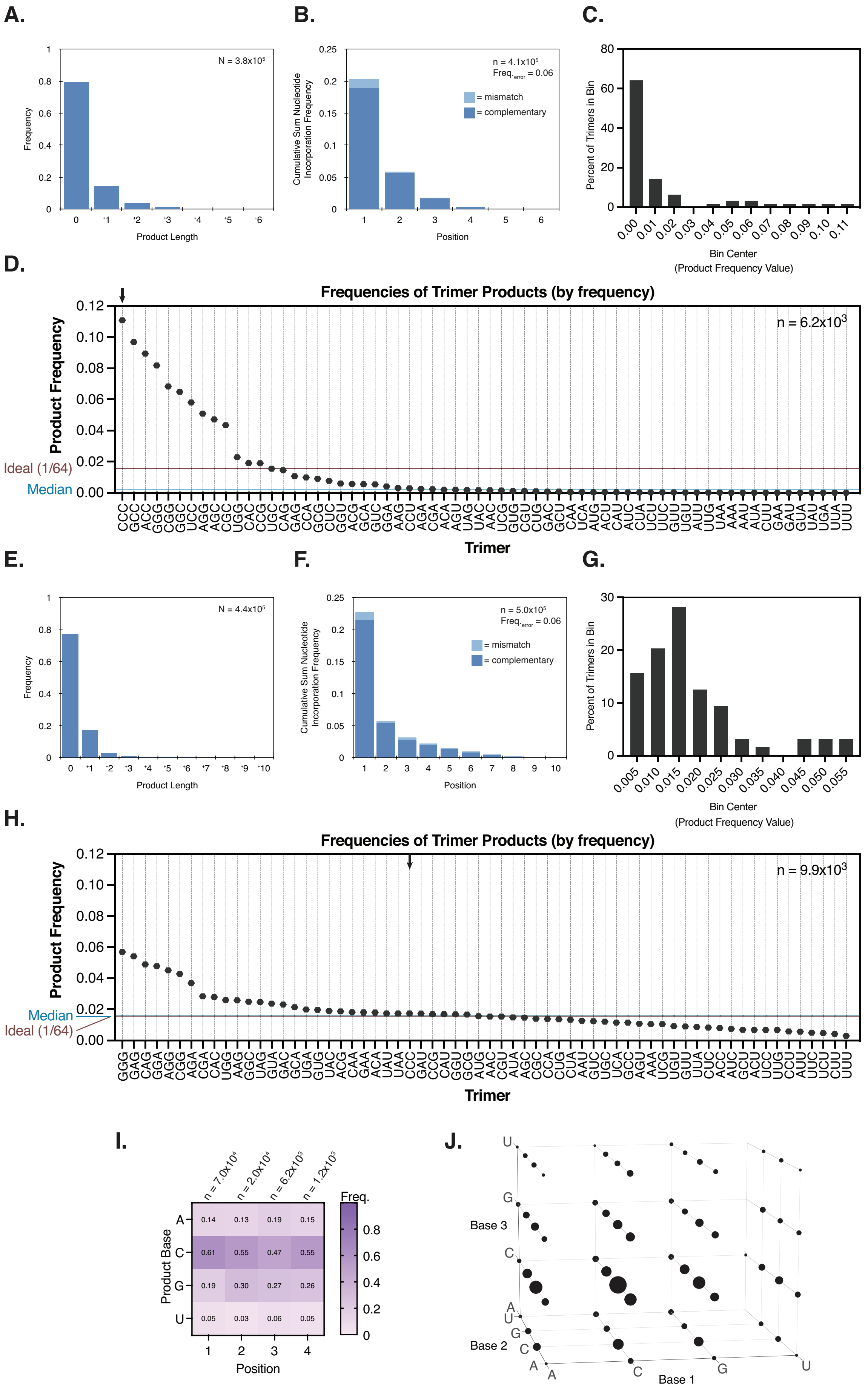

A.
N = 3.8x10^5
Frequency
Product Length
B.
n = 4.1x10^5
Freq.error = 0.06
= mismatch
= complementary
Cumulative Sum Nucleotide Incorporation Frequency
Position
C.
Percent of Trimers in Bin
Bin Center
(Product Frequency Value)
D.
Frequencies of Trimer Products (by frequency)
n = 6.2x10^3
Product Frequency
Ideal (1/64)
Median
Trimer
E.
N = 4.4x10^5
Frequency
Product Length
F.
n = 5.0x10^5
Freq.error = 0.06
= mismatch
= complementary
Cumulative Sum Nucleotide Incorporation Frequency
Position
G.
Percent of Trimers in Bin
Bin Center
(Product Frequency Value)
H.
Frequencies of Trimer Products (by frequency)
n = 9.9x10^3
Product Frequency
Median
Ideal (1/64)
Trimer
I.
n = 7.0x10^4
n = 2.0x10^4
n = 6.2x10^3
n = 1.2x10^3
Freq.
Product Base
Position
J.
Base 3
Base 2
Base 1

**Supplementary Figure 3. Additional comparisons of primer extension without and with added oligonucleotides. A-D.** Product sequence features of primer extension with 20 mM pre-activated 2AIrN and MeNC activation chemistry to drive bridged dinucleotide formation. **A.** The length distribution of all products, normalized to all lengths including 0 (unextended). Note that yields are relatively low. **B.** The cumulative sum frequency distribution of correct and incorrect nucleotide incorporations normalized to each position. The frequency of incorrect incorporations is 0.063. **C.** The distribution of complementary products at least three nucleotides long is skewed. Many trimer products are poorly represented. **D.** Quantification of complementary product trimer frequencies, ordered by frequency (same data as in Figure 2C). Note the high frequencies of G/C-rich trimers. The arrow indicates the highest frequency trimer: CCC. **E-H.** Product sequence features of primer extension with 20 mM rN + 1 mM oligo mix ($pN_{2\text{-}16}$) + *in situ* MeNC activation. **E.** The length distribution of all products, normalized to all lengths including 0 (unextended). The yields are similar to those without the oligo mix (A). Note the low-frequency tail of ligation products up to +6, especially visible in the **F.** cumulative sum frequency distribution of correct and incorrect nucleotide incorporations normalized to each position. The frequency of incorrect incorporations is 0.063, as for (A-D). **G.** The distribution of complementary products at least three nucleotides long is more normal than without the oligo mix (compare to (C)). **H.** Quantification of complementary product trimer frequencies, ordered by frequency (same data as in Figure 2F). **I-J.** Product sequence features of primer extension with an *in situ* activated mixture of 20 mM rN + 250 µM $pN_3$. Under experimental conditions identical to those in Figure 2D-F and Supplementary Figure 3E-H, but using only additional trimer oligonucleotides instead of the oligo mix, does not yield unbiased sequence products. **I.** The position-normalized frequency distribution of products remains biased to rG and rC, and **J.** the distribution of trimer products is more uniform than without the added trimer oligonucleotides, but still more biased than with the added oligo mix (compare to Figure 2D and 2E). The error frequency is 0.15.

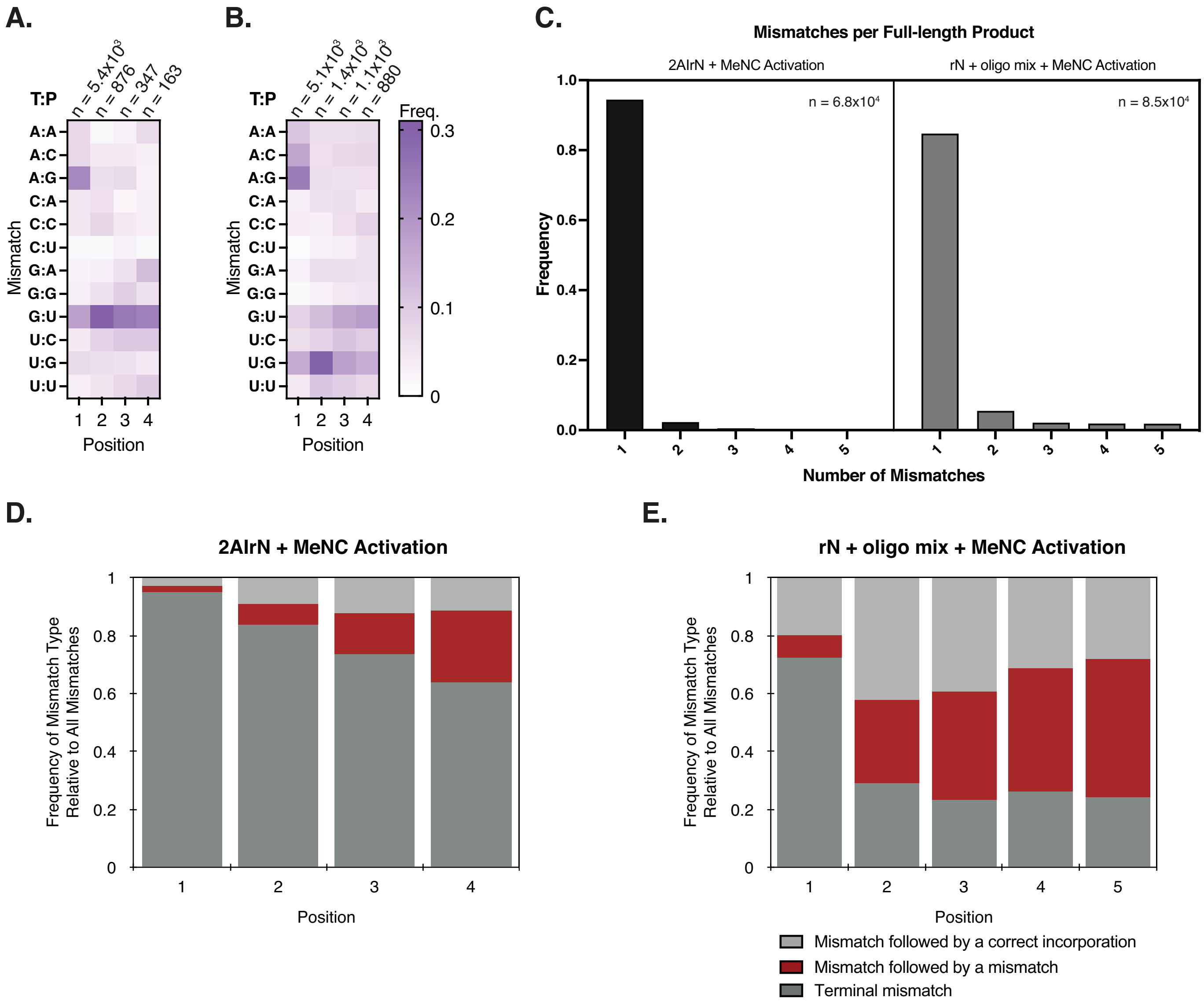

**Supplementary Figure 4. The inclusion of the oligo mix during primer extension alters the distribution of mismatches. A, C, & D.** Sequence features of products with mismatches from primer extension with 20 mM pre-activated 2AIrN and MeNC activation chemistry to drive bridged dinucleotide formation. **B, C, & E.** Sequence features of products with mismatches from primer extension with 20 mM rN + 1 mM oligo mix ($pN_{2\text{-}16}$) + *in situ* MeNC activation. **A.** The distribution of mismatches (listed as **T**emplate:**P**roduct) exhibits the known pattern for reactions with all four nucleotides and a random template (1). **B.** The distribution of mismatches is different in the reaction that includes the oligo mix. G:U and U:G wobble base pairs are more highly represented. **C.** Mismatches are typically terminal in reactions with just 2AIrN (4). Therefore, most polymerization products with a mismatch contain only one mismatch (left panel). Products harboring multiple mismatches are more common with the inclusion of the oligo mix (right panel). **D.** Mismatches can be grouped into three categories: 1. terminal (dark grey), 2. followed by a mismatch (red), or 3. followed by a correct incorporation (light grey). Consistent with (**C**), most mismatches are terminal in reactions with just 2AIrN. **E.** Non-terminal mismatches are more common at all positions in the reaction that includes the oligo mix. This is consistent with a contribution from oligonucleotide ligations.

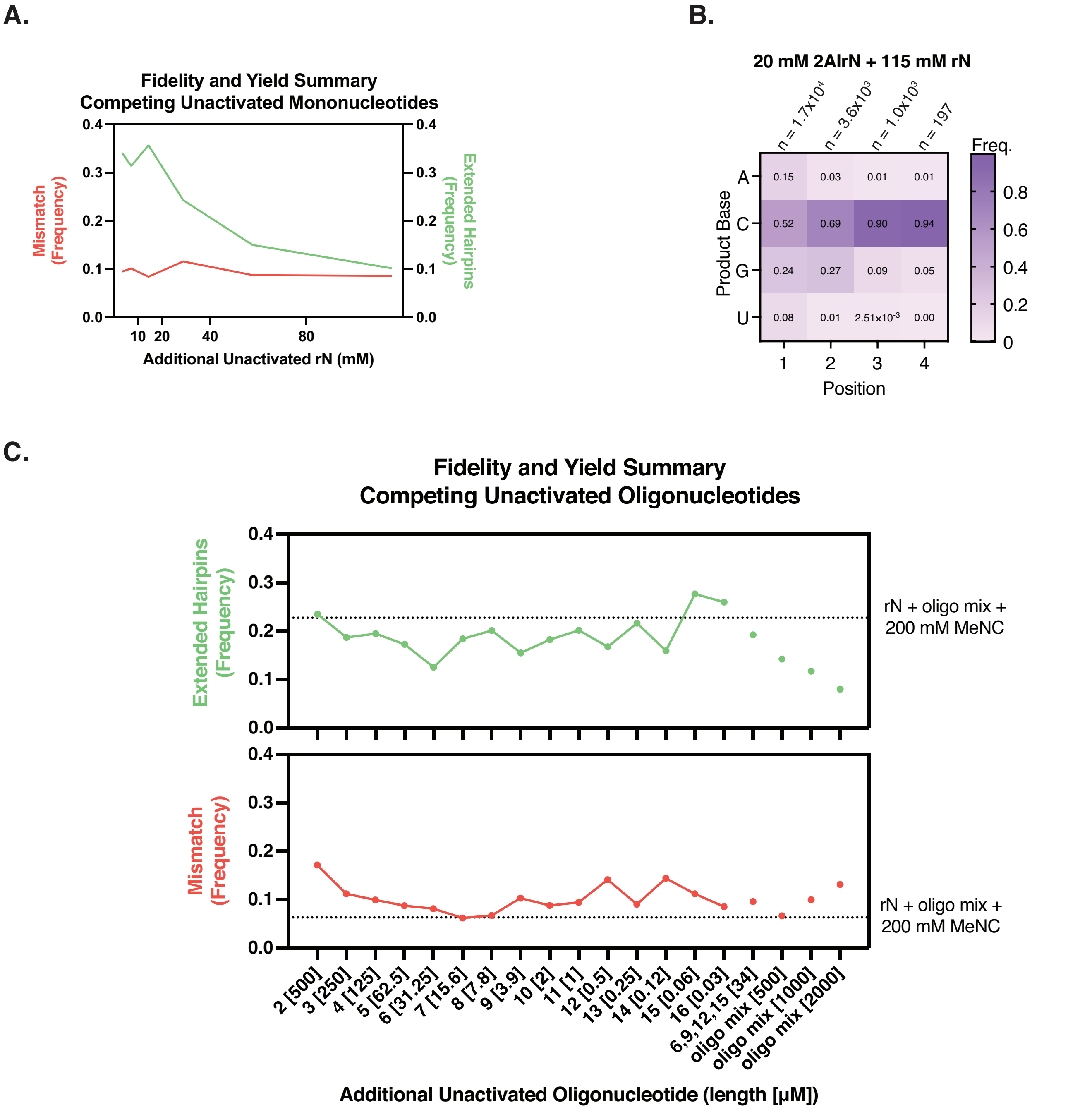

**Supplementary Figure 5. Competing unactivated mononucleotides decrease product yields, and competing unactivated oligonucleotides increase mismatch frequencies. A.** In a primer extension reaction with 20 mM 2AIrN, the addition of unactivated nucleotides at increasing concentrations progressively reduces product yields. Product yields (green) are quantified as the frequency of hairpin primers extended by at least one nucleotide. There is no significant effect on mismatch frequencies (red) or **B.** complementary product sequence makeup (compare to Figure 2A). **C.** In a primer extension reaction with 20 mM 2AIrN, the addition of unactivated sets of competitor oligonucleotides of individual lengths has an inconsistent but relatively minor effect on product yields (green). The addition of the oligo mix (1 mM) measurably reduces yields relative to the rN + oligo mix + MeNC experiment (dotted line; see Supplementary Figure 3F). The addition of any individual oligonucleotide except $pN_7$, a subset of longer oligonucleotides ($pN_6$, $pN_9$, $pN_{12}$, + $pN_{15}$), the oligo mix at the standard concentration (1 mM), or the oligo mix at twice the standard concentration (2 mM) all increase mismatch frequencies (red) relative to the rN + oligo mix + MeNC experiment (dotted line; see Figure 2D-F and Supplementary Figure 3E-H). The addition of the oligo mix at half the standard concentration (0.5 mM) does not significantly increase mismatches.

One possible explanation for these mismatch trends is that oligonucleotides bound to the template but containing a mismatch are likely to harbor that mismatch at a terminus (see Supplementary Figure 9). If such a 5′ mismatch in the oligo sandwiches an activated mononucleotide or bridged dinucleotide against the primer then it may potentiate the incorporation of a mismatch by distorting the local helical structure. In these datasets, oligonucleotides longer than 7 are statistically very likely to harbor a mismatch when bound to the template (Supplementary Figure 10A). The inclusion of various mixtures of oligonucleotides in the primer extension reaction results in somewhat lower mismatch frequencies than the inclusion of many of the individual oligonucleotides. This could be explained by more frequent and stable off-template interactions among oligonucleotides in a mixture of lengths, thereby decreasing the relative availability of any individual oligonucleotide to interfere in the reaction. Additionally, off-template oligonucleotides may titrate bridged dinucleotides relative to activated mononucleotides. Additional work will be required to explore these hypotheses.

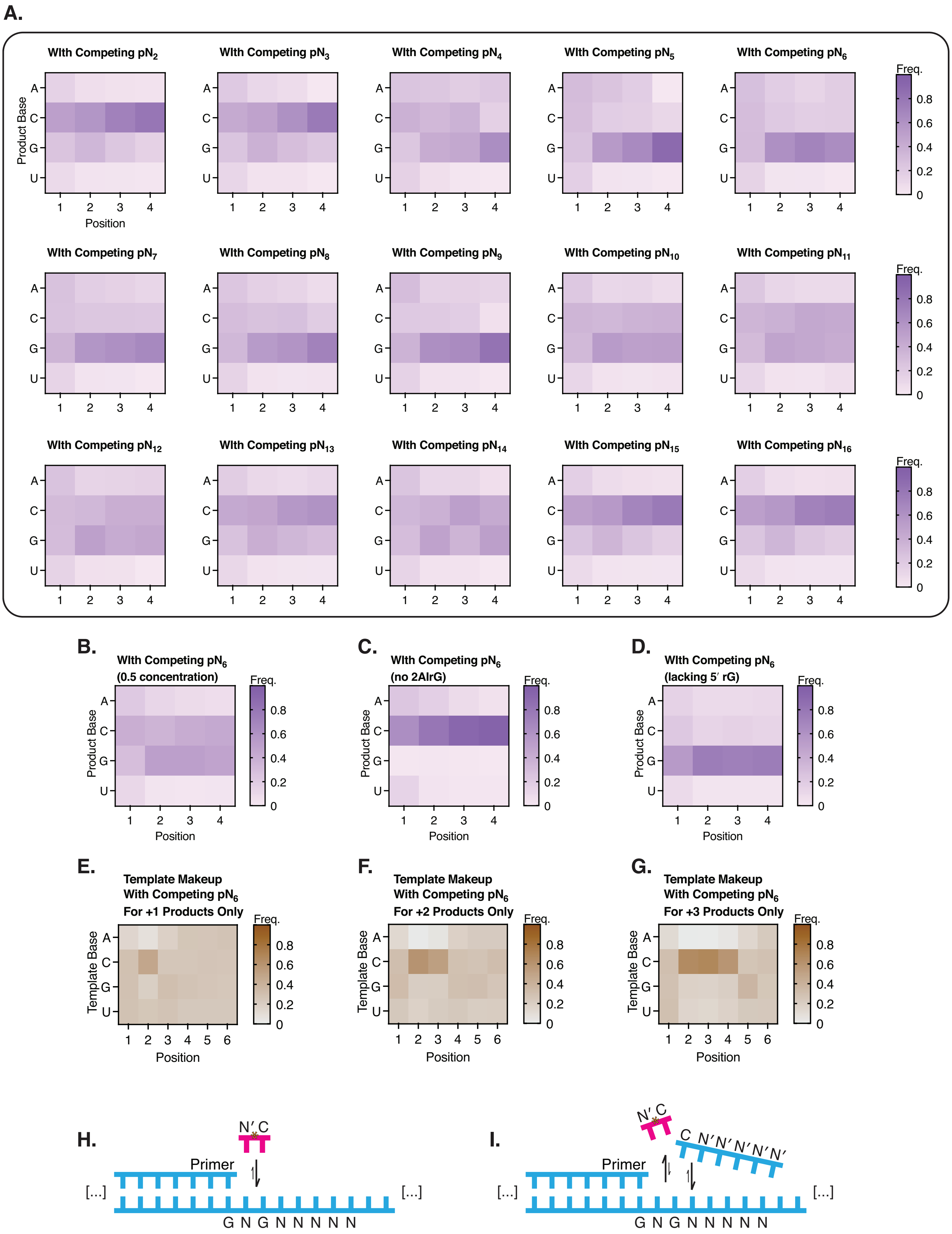
A.
With Competing $pN_2$
With Competing $pN_3$
With Competing $pN_4$
With Competing $pN_5$
With Competing $pN_6$
With Competing $pN_7$
With Competing $pN_8$
With Competing $pN_9$
With Competing $pN_{10}$
With Competing $pN_{11}$
With Competing $pN_{12}$
With Competing $pN_{13}$
With Competing $pN_{14}$
With Competing $pN_{15}$
With Competing $pN_{16}$
Product Base
A
C
G
U
Position
1 2 3 4
Freq.
0 0.2 0.4 0.6 0.8
B.
With Competing $pN_6$ (0.5 concentration)
C.
With Competing $pN_6$ (no 2AIrG)
D.
With Competing $pN_6$ (lacking 5′ rG)
E.
Template Makeup With Competing $pN_6$ For +1 Products Only
F.
Template Makeup With Competing $pN_6$ For +2 Products Only
G.
Template Makeup With Competing $pN_6$ For +3 Products Only
Template Base
1 2 3 4 5 6
H.
N′ C
Primer
[...]
[...]
G N G N N N N N
I.
N′ C
C N′ N′ N′ N′ N′
Primer
[...]
[...]
G N G N N N N N

**Supplementary Figure 6. Competing unactivated oligonucleotides change the complementary product distributions. A.** The position-normalized frequency distributions of complementary products for primer extension reactions with 20 mM 2AIrN incubated for 24 hours + the set of unactivated oligonucleotide of the indicated length (same data as Supplementary Figure 5C; 500 µM $pN_2$; 250 µM $pN_3$; 125 µM $pN_4$; 62.5 µM $pN_5$; 31.3 µM $pN_6$; 15.6 µM $pN_7$; 7.8 µM $pN_8$; 3.9 µM $pN_9$; 2 µM $pN_{10}$; 0.98 µM $pN_{11}$; 0.49 µM $pN_{12}$; 0.24 µM $pN_{13}$; 0.12 µM $pN_{14}$; 0.06 µM $pN_{15}$; or 0.03 µM $pN_{16}$). Products are rC-rich in the reaction with just 2AIrN (see Figure 2A). The addition of $pN_2$ or $pN_3$ at these concentrations does not change the product distributions, which remain rC-rich as in the absence of added oligos. $pN_4$ and longer unactivated competing oligonucleotides up to $pN_9$ induce a rG-rich pattern of products. Unactivated competing oligos of lengths $pN_{10}$-$pN_{14}$ induce an intermediate rC/G-rich pattern of products. Finally, oligos $pN_{15}$-$pN_{16}$ are ineffective competitors, and the complementary product distribution reverts to rC-rich. **B.** The product distribution for a reaction with 20 mM 2AIrN incubated for 24 hours + $pN_6$ at half the concentration used in (A) (15.6 µM instead of 31.3 µM). The product distribution pattern is less rG-rich when the concentration of $pN_6$ is reduced. This result is consistent with the hypothesis that the change in product distributions is driven by competition for template binding sites. **C.** The product distribution for a reaction with 20 mM 2AIrA, U, and C (*no* 2AIrG) incubated for 24 hours + competing unactivated $pN_6$. Even with competing $pN_6$, which demonstrably inhibits rC products and thereby favors rG products, the rC-rich product distribution can be recovered by excluding 2AIrG as a reactant. This indicates that the competition for rG templates by the oligonucleotides is dynamic. That is, $pN_6$ oligos are not permanently bound to the template. **D.** The product distribution for a reaction with 20 mM 2AIrN incubated for 24 hours + a special $pN_6$ oligonucleotide lacking 5′-rG (p(A,U,C)NNNNN). The rG-rich product distribution is retained with this oligo. This indicates that the rG-rich products are not a consequence of selection for rC-rich templates by passive oligonucleotide sandwiching of either mononucleotides or bridged dinucleotides. (For example, an oligo with a 5′-rG could sandwich an N*N bridged dinucleotide bound to template positions 1 and 2. That bridged dinucleotide would then react with the primer, and thereby enrich the frequency of rC in template position 3 among templates that are extended by at least one nucleotide. If this were a dominant factor then excluding the 5′-rG in $pN_6$ should reduce rG-rich products, but it does not.) **E-G.** The position-normalized frequency distributions of *templating bases* for complementary products of lengths +1, +2, or +3 for reactions with 20 mM 2AIrN incubated for 24 hours + $pN_6$ (same experiment as in [A]). **E.** Among products that extended to +1, there is a strong overrepresentation of rC templates in position 2. This is consistent with efficient reactivity by N*G bridged dinucleotides, instead of N*C bridged dinucleotides, in the presence of competing $pN_6$. The same pattern holds for **F.** products that extended to +2 and **G.** +3. We conclude that the rG-rich product distribution observed with competing unactivated oligonucleotides of lengths $pN_{5\text{-}9}$ is driven by the relatively high reactivity of N*G bridged dinucleotides in the presence of competing oligos. **H-I.** One possible mechanism to explain the observed results. **H.** N*C bridged dinucleotides are highly reactive in the absence of competing unactivated oligonucleotides but **I.** their ability to bind the template is inhibited by rC-rich oligonucleotides.

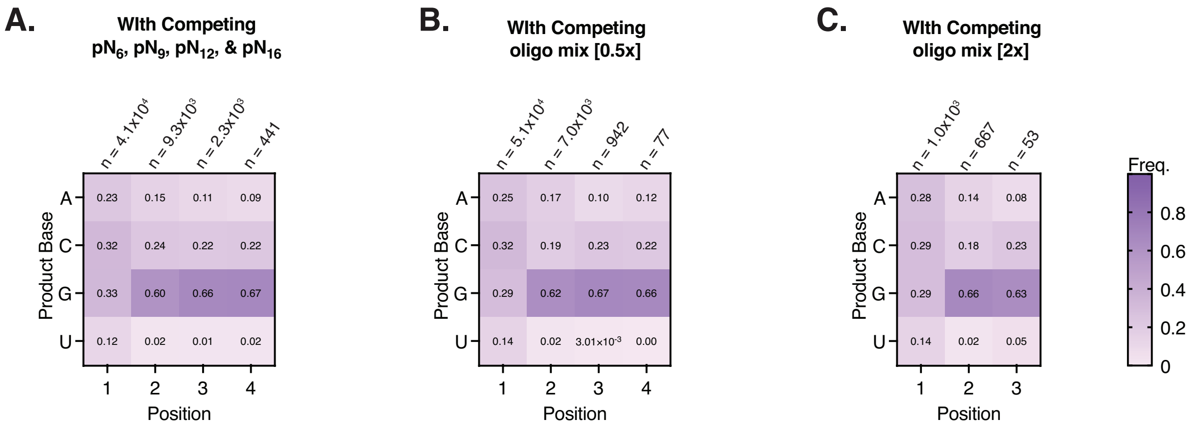

**Supplementary Figure 7. Competing unactivated oligonucleotide mixtures bias the complementary product distribution to rG-rich. A.** The position-normalized frequency distribution of complementary products for a primer extension reaction with 20 mM 2AIrN incubated for 24 hours + (31.3 µM $pN_6$, 3.9 µM $pN_9$, 0.49 µM $pN_{12}$, and 0.03 µM $pN_{16}$). **B.** The product distribution for a reaction with 20 mM 2AIrN incubated for 24 hours + 0.5 mM oligo mix. **C.** The product distribution for a reaction with 20 mM 2AIrN incubated for 24 hours + 2 mM oligo mix.

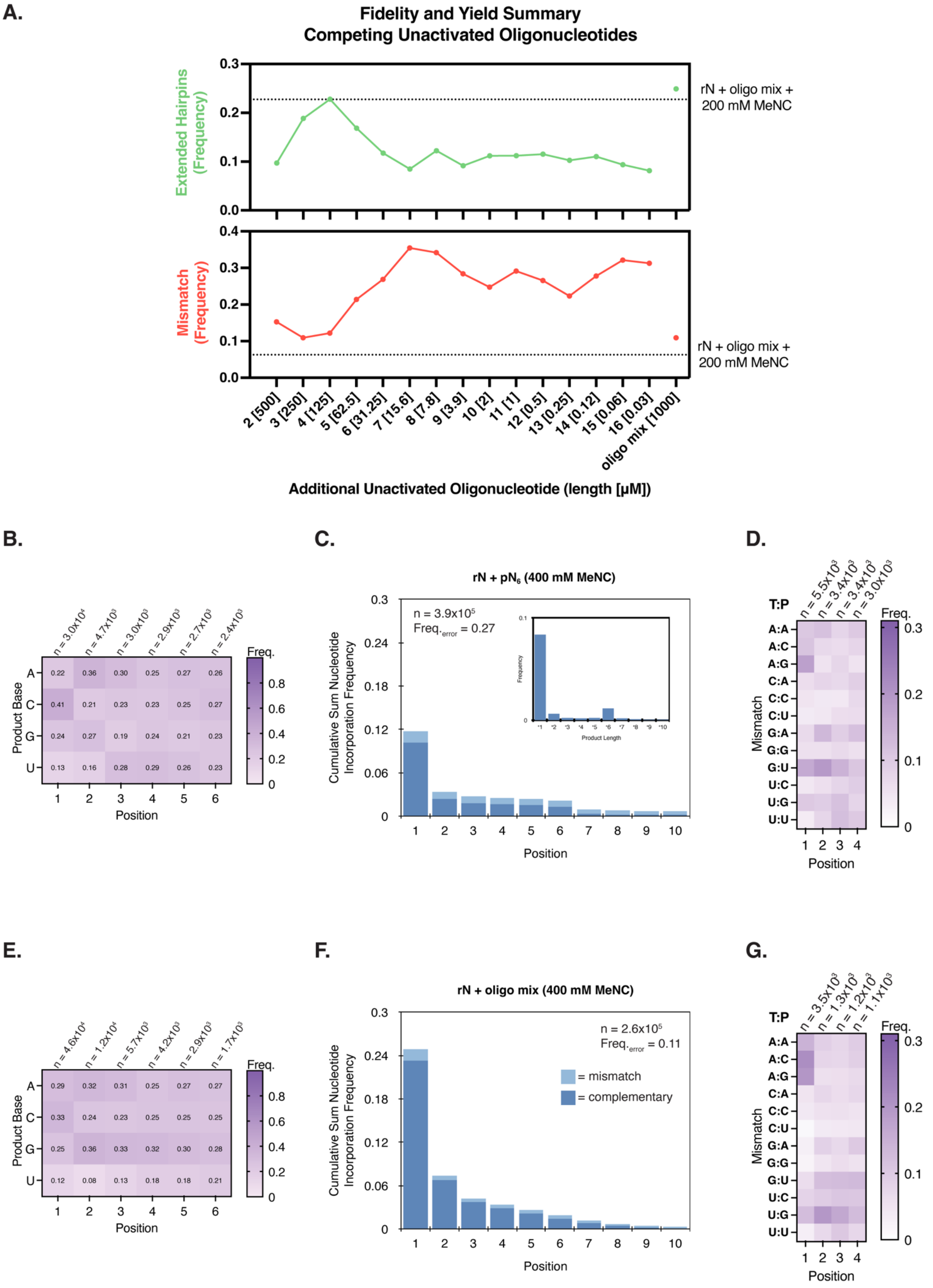

A.
Fidelity and Yield Summary
Competing Unactivated Oligonucleotides
Extended Hairpins (Frequency)
Mismatch (Frequency)
rN + oligo mix + 200 mM MeNC
rN + oligo mix + 200 mM MeNC
2 [500]
3 [250]
4 [125]
5 [62.5]
6 [31.25]
7 [15.6]
8 [7.8]
9 [3.9]
10 [2]
11 [1]
12 [0.5]
13 [0.25]
14 [0.12]
15 [0.06]
16 [0.03]
oligo mix [1000]
Additional Unactivated Oligonucleotide (length [μM])
B.
n = 3.0x10⁴
n = 4.7x10³
n = 3.0x10³
n = 2.9x10³
n = 2.7x10³
n = 2.4x10³
Product Base
A 0.22 0.36 0.30 0.25 0.27 0.26
C 0.41 0.21 0.23 0.23 0.25 0.27
G 0.24 0.27 0.19 0.24 0.21 0.23
U 0.13 0.16 0.28 0.29 0.26 0.23
Position
Freq.
C.
rN + pN₆ (400 mM MeNC)
n = 3.9x10⁵
Freq.error = 0.27
Cumulative Sum Nucleotide Incorporation Frequency
Frequency
Product Length
Position
D.
T:P
n = 5.5x10³
n = 3.4x10³
n = 3.4x10³
n = 3.0x10³
A:A
A:C
A:G
C:A
C:C
C:U
G:A
G:G
G:U
U:C
U:G
U:U
Mismatch
Position
Freq.
E.
n = 4.6x10⁴
n = 1.2x10⁴
n = 5.7x10³
n = 4.2x10³
n = 2.9x10³
n = 1.7x10³
Product Base
A 0.29 0.32 0.31 0.25 0.27 0.27
C 0.33 0.24 0.23 0.25 0.25 0.25
G 0.25 0.36 0.33 0.32 0.30 0.28
U 0.12 0.08 0.13 0.18 0.18 0.21
Position
Freq.
F.
rN + oligo mix (400 mM MeNC)
n = 2.6x10⁵
Freq.error = 0.11
= mismatch
= complementary
Cumulative Sum Nucleotide Incorporation Frequency
Position
G.
T:P
n = 3.5x10³
n = 1.3x10³
n = 1.2x10³
n = 1.1x10³
A:A
A:C
A:G
C:A
C:C
C:U
G:A
G:G
G:U
U:C
U:G
U:U
Mismatch
Position
Freq.

**Supplementary Figure 8. High-concentration MeNC activation increases error frequencies. A-G.** Reactions with initially unactivated 20 mM rN + the indicated oligonucleotide or oligo mix subjected to eutectic 400 mM MeNC activation followed by solution-phase 400 mM MeNC activation (instead of the standard 200 mM MeNC, as in Figure 2D-F and Supplementary Figure 3E-H). **A.** The inclusion of any set of oligonucleotides of one length in the primer extension reaction reduces yields (green) and increases mismatches (red) relative to the standard reaction with the oligo mix and 200 mM MeNC (dotted lines). The inclusion of the oligo mix and 400 mM MeNC (last data point) shows higher yields and lower error frequencies than all the reactions with a set of oligonucleotide of one length and 400 mM MeNC. **B-D.** Product sequence features for the representative example of 20 mM rN + 31.3 µM $pN_6$ + 400 mM MeNC activation. **B.** The position-normalized frequency distribution of complementary products is relatively unbiased. **C.** The cumulative sum frequency distribution of correct and incorrect nucleotide incorporations normalized to each position shows a strong contribution to errors from ligations. Note the prominent 6-nucleotide long tail of product from $pN_6$ ligation. The frequency of incorrect incorporations is 0.27. The inset shows the product length distribution from the same experiment, to highlight the contribution of $pN_6$ ligation: products of length +6 are clearly overrepresented. **D.** The distribution of mismatches (listed as **T**emplate:**P**roduct) shows a pattern intermediate between that found in the primer extension reaction with just 2AIrN and with oligonucleotide ligations (Supplementary Figure 4A & B and Supplementary Figure 10C). **E-G.** Product sequence features for a reaction with 20 mM rN + 1 mM oligo mix + 400 mM MeNC activation. **E.** With the inclusion of the oligo mix, the product distribution is unbiased, and **F.** yields increase and errors decrease relative to reactions with just a set of oligonucleotides of one length. The mismatch frequency of 0.11 is lower than with any individual oligonucleotide, but higher than that with 200 mM MeNC (error frequency of 0.063, Supplementary Figure 3F). **G.** The distribution of mismatches is again intermediate between that found in the primer extension reaction with just 2AIrN and with oligonucleotide ligations. It is similar to the mismatch distribution for the same reaction but with 200 mM MeNC instead of 400 mM MeNC (Supplementary Figure 4B).

| Entry | Oligo | Activating Group | Activation Chemistry | Frequency of Oligo-length Product |
|---|---|---|---|---|
| **1** | 500 µM 2AIpCGG | 2AI | Pre-activated and purified oligo | 0.28 |
| **2** | 250 µM pCGG | 50 mM 2AI | Solution-phase 200 mM MeNC | 0.0012 |
| **3** | 250 µM pCGG | 100 mM 2AI | Solution-phase 200 mM MeNC | 0.00082 |
| **4** | 250 µM pCGG | 200 mM 2AI | Solution-phase 200 mM MeNC | 0.0017 |
| **5** | 250 µM pCGG | 20 mM 2AI | Eutectic 200 mM MeNC activation followed by solution-phase 200 mM MeNC | 0.0028 |
| **6** | 250 µM $pN_3$ | 20 mM 2AI | Eutectic 200 mM MeNC activation followed by solution-phase 200 mM MeNC | 0.0024 |
| **7** | 500 µM $pN_2$ | 200 mM 2MI | Solution-phase 200 mM MeNC | 0.00084 |
| **8** | 250 µM $pN_3$ | 200 mM 2MI | Solution-phase 200 mM MeNC | 0.022 |
| **9** | 250 µM $pN_3$ | 200 mM 1MI | Solution-phase 400 mM MeNC | 0.025 |

**Supplementary Table 1, Test Ligation Experiments**

**Supplementary Table 1. *In situ* activation of oligonucleotides is inefficient. Entry 1** shows a ligation frequency of 0.28 for a pre-activated 2AIpCGG trimer on a GCC template after a 24 hour incubation (3). **Entries 2-6.** Attempts to generate comparable ligation yields by activating the same pCGG or $pN_3$ *in situ* with 2AI failed and resulted in statistically negligible yields. Because ligation is efficient once the oligo is activated (see Entry 1), we conclude that the initial activation of just oligonucleotides with MeNC is inefficient. **Entries 7-8.** 2-methylimidazole (2MI) improves product yields. This is probably due to the higher reactivity of this leaving group rather than more efficient initial activation. **Entry 9.** The best test result for a ligation-only reaction was with 1-methylimidazole (1MI), 400 mM MeNC solution-phase activation, and a 48 hour incubation (unlike all other entries, which were for 24 hours).

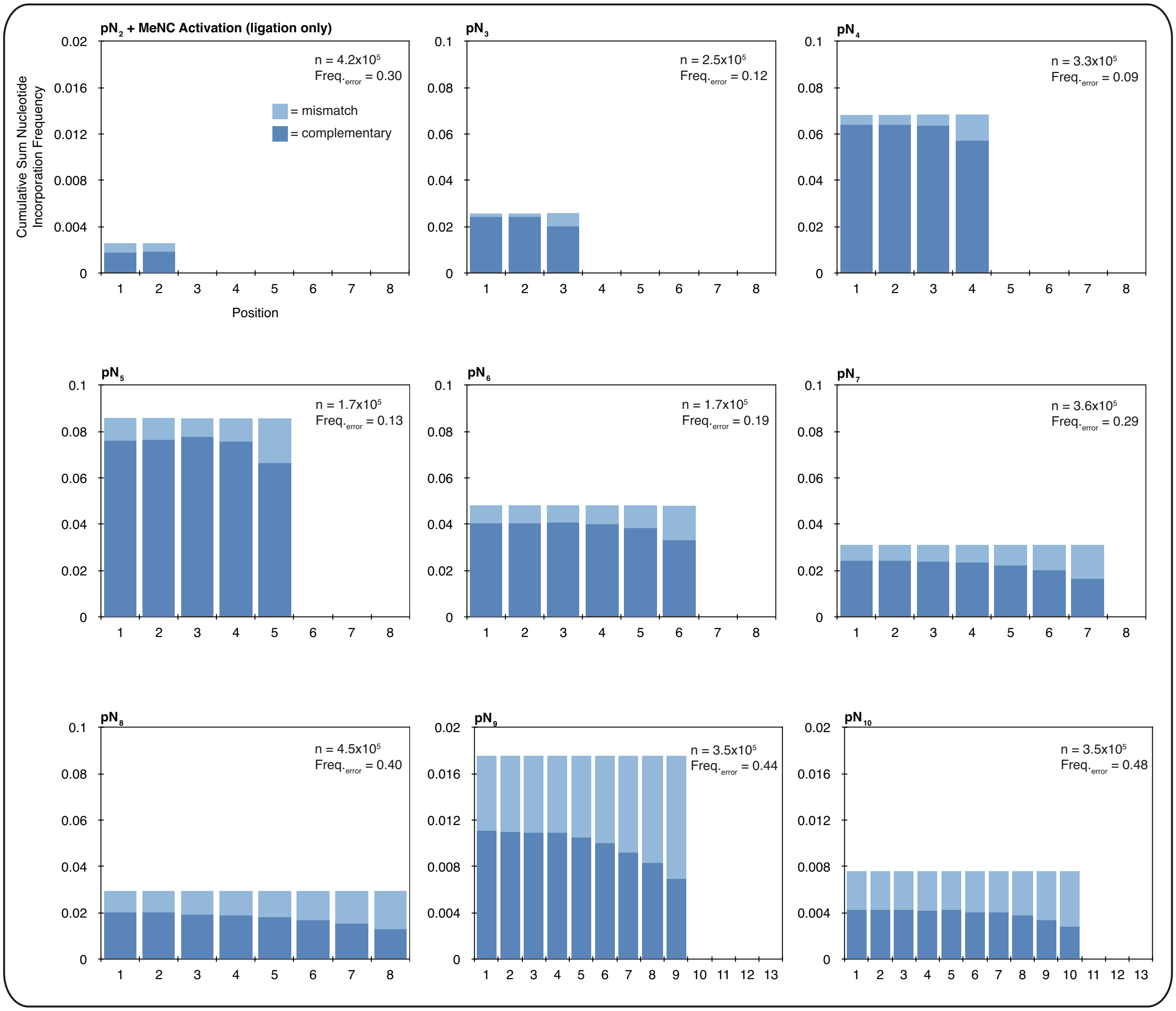

**Supplementary Figure 9. Oligonucleotide ligation-only reactions can be error-prone, with the highest frequency of mismatches at the 3′-terminus of ligation products.** The cumulative sum frequency distributions of correct and incorrect nucleotide incorporations normalized to each position for ligation-only reactions with 200 mM 1-methylimidazole (1MI), 400 mM MeNC solution-phase activation, and the indicated set of oligonucleotides of one length incubated for 48 hours (500 µM $pN_2$; 250 µM $pN_3$; 125 µM $pN_4$; 62.5 µM $pN_5$; 31.3 µM $pN_6$; 15.6 µM $pN_7$; 7.8 µM $pN_8$; 3.9 µM $pN_9$; 2 µM $pN_{10}$). The data were filtered for oligo-length products to reduce noise. The yields are relatively low throughout (compare to Supplementary Figure 3), and the mismatch frequencies are relatively high. The highest mismatch frequency is consistently at the 3′-most nucleotide, except with $pN_2$. (Note the y-axis scale change between $pN_2$ and $pN_3$, and between $pN_8$ and $pN_9$. Reactions with $pN_{11-16}$ showed yields too low for statistically clear interpretation.)

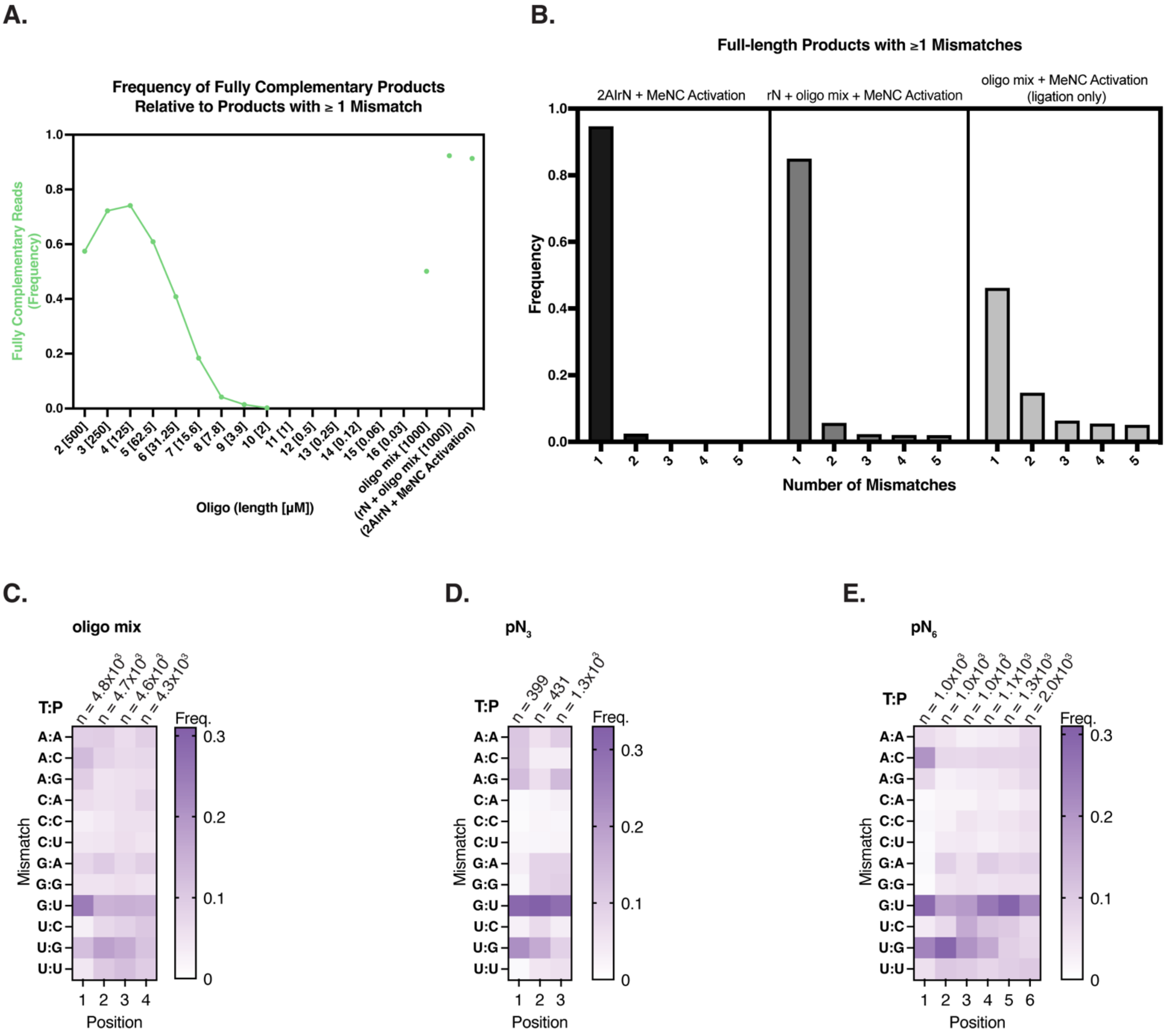

**Supplementary Figure 10. Oligonucleotide ligation-only reactions show a pattern of mismatches that is distinct from that observed for reactions with 2AIrN. A-E.** Mismatch features for ligation-only reactions with 200 mM 1-methylimidazole (1MI), 400 mM MeNC solution-phase activation, and the indicated set of oligonucleotides of one length or oligo mix incubated for 48 hours. **A.** The frequency of products that are fully complementary as a function of ligated oligo length. Note the sharp drop in fully complementary ligation products for oligos ≥ $pN_5$. Primer extension ligation reactions with $pN_{11-16}$ showed yields too low for statistically clear interpretation. The ligation of the oligo mix resulted in a frequency of only 0.50 fully complementary products, compared to 0.92 for the rN + oligo mix + MeNC reaction, and 0.91 for the reaction with pre-activated 2AIrN and MeNC activation. **B.** Among products with at least one mismatch from the primer extension reaction with 2AIrN, most contain just one mismatch (left panel). Products with multiple mismatches are slightly more common in the reaction with rN + oligo mix + MeNC activation (middle panel) (same data as in Supplementary Figure 4C, shown here for comparison). However, for the ligation-only experiment with the oligo mix, more than half of the products with at least one mismatch contain ≥ 2 mismatches. Note how the middle panel is intermediate between the case with just 2AIrN and the case with just ligation. The distributions of mismatches (listed as **T**emplate:**P**roduct) for **C.** ligation of the oligo mix, **D.** ligation of $pN_3$, and **E.** ligation of $pN_6$ are similar, and distinct from the case with just 2AIrN reactants (see Supplementary Figure 4A).

**A.**

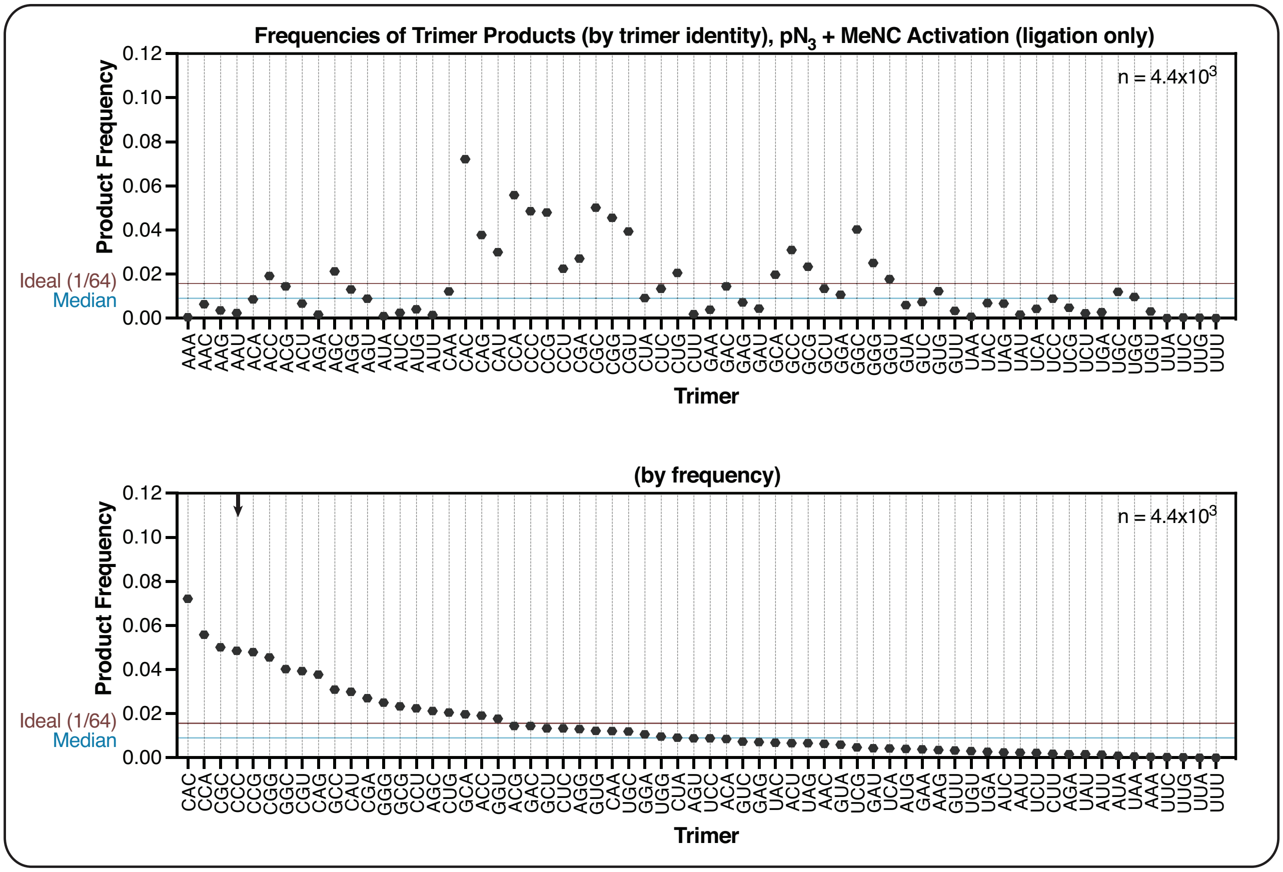

Frequencies of Trimer Products (by trimer identity), $pN_3$ + MeNC Activation (ligation only)
Product Frequency
Trimer
Ideal (1/64)
Median
$n = 4.4x10^3$
(by frequency)
Product Frequency
Trimer
Ideal (1/64)
Median
$n = 4.4x10^3$

**B.**

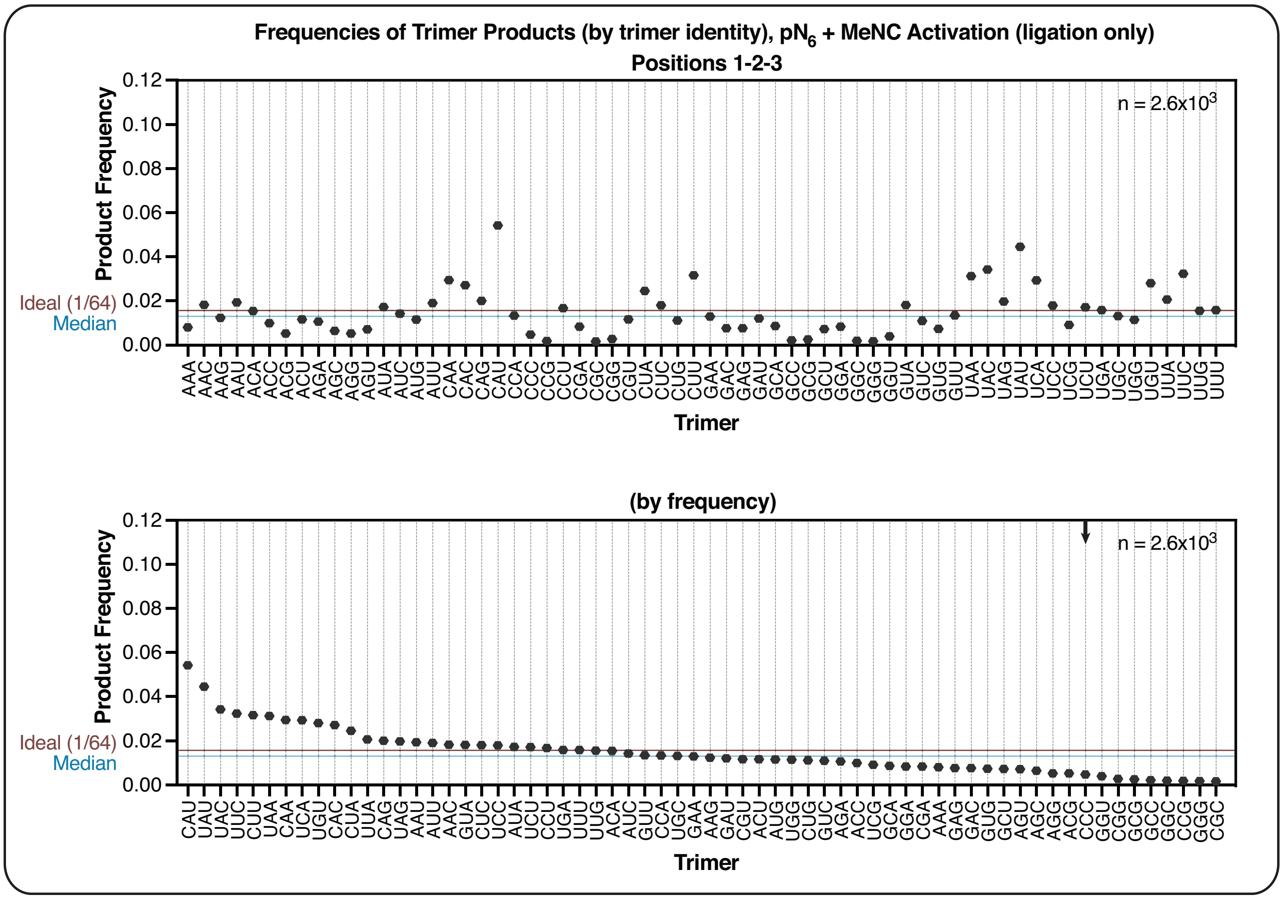

Frequencies of Trimer Products (by trimer identity), $pN_6$ + MeNC Activation (ligation only)
Positions 1-2-3
Product Frequency
Trimer
Ideal (1/64)
Median
$n = 2.6x10^3$
(by frequency)
Product Frequency
Trimer
Ideal (1/64)
Median
$n = 2.6x10^3$

C.

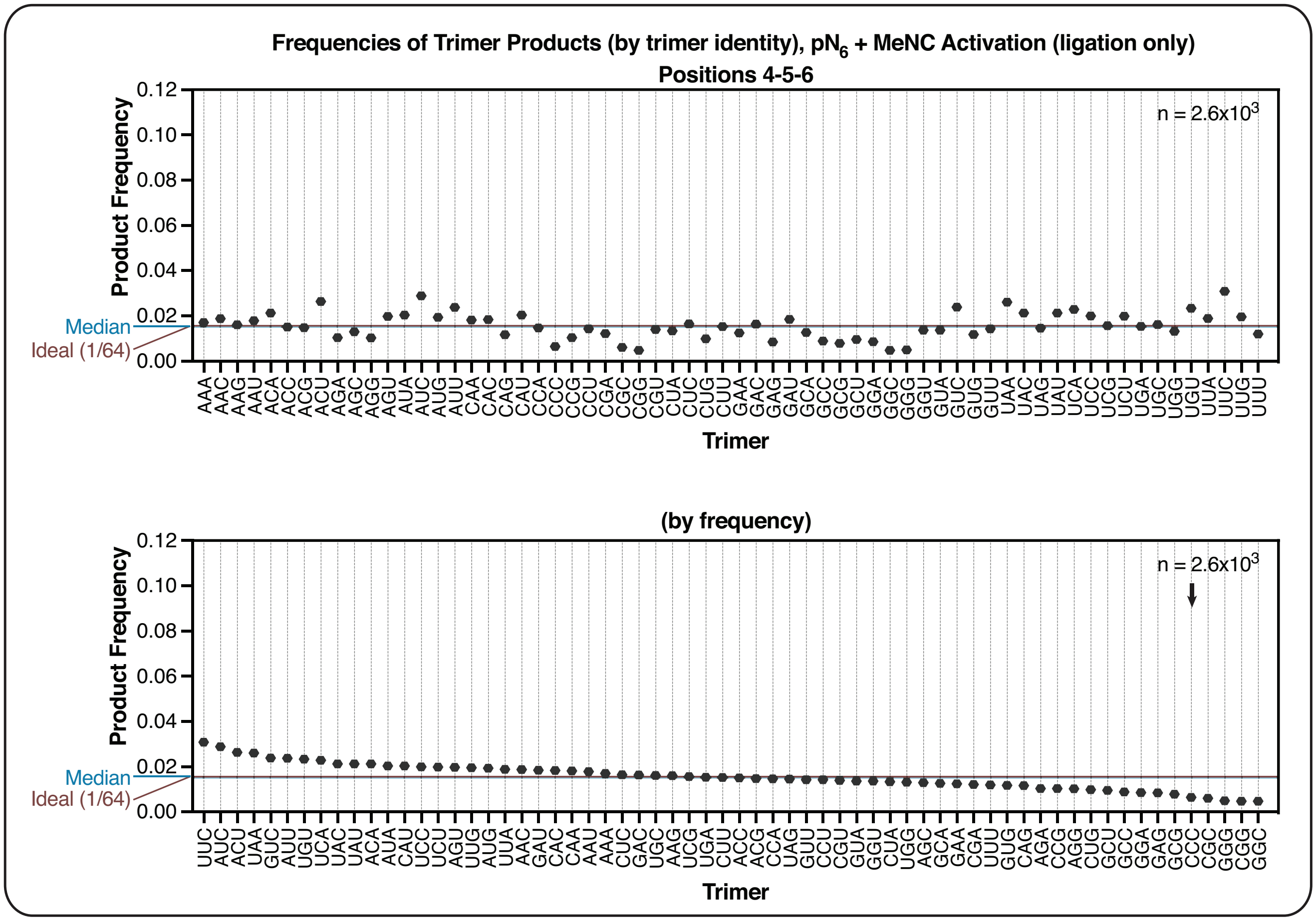

**Supplementary Figure 11. Ligation-only reactions exhibit complementary product sequence biases as a function of oligonucleotide length. A-C.** Quantification of product trimer frequencies for templated ligation with $pN_3$ or $pN_6$, 200 mM 1-methylimidazole (1MI), and 400 mM MeNC solution-phase activation. **A.** $pN_3$ ligation yields a biased distribution of products, comparable to that measured for the primer extension reaction with 2AIrN (compare with Figure 2C and Supplementary Figure 3D). rC- and rG-rich products are overrepresented and rA- and rU-rich products are underrepresented. The CCC trimer, which has the highest frequency in reactions with 2AIrN, is indicated by the arrow throughout. The median frequency is 0.009. The ideal frequency for each trimer is 1/64 = 0.016. **B.** $pN_6$ ligation yields a distinct distribution of product trimers at positions 1-2-3 that is more uniform than for $pN_3$ ligation. The trimer products are biased *against* rC- and rG-rich products, with strong representation of rA and rU. The median product trimer frequency is 0.013. **C.** The distribution of product trimer frequencies from $pN_6$ ligation at positions 4-5-6. The median frequency is 0.015.

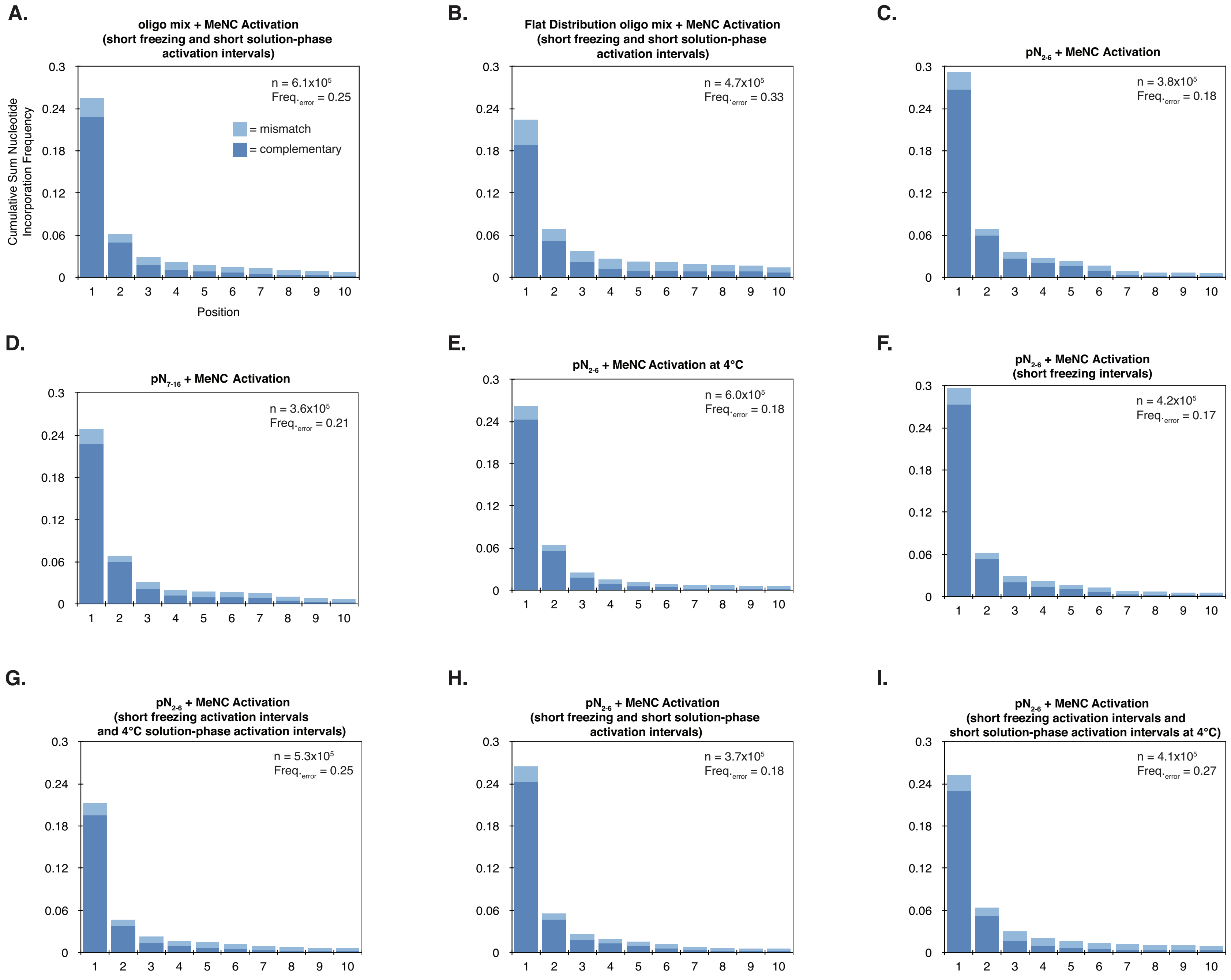
A.
oligo mix + MeNC Activation (short freezing and short solution-phase activation intervals)
n = 6.1x10^5
Freq.error = 0.25
= mismatch
= complementary
Cumulative Sum Nucleotide Incorporation Frequency
Position
B.
Flat Distribution oligo mix + MeNC Activation (short freezing and short solution-phase activation intervals)
n = 4.7x10^5
Freq.error = 0.33
C.
pN2-6 + MeNC Activation
n = 3.8x10^5
Freq.error = 0.18
D.
pN7-16 + MeNC Activation
n = 3.6x10^5
Freq.error = 0.21
E.
pN2-6 + MeNC Activation at 4°C
n = 6.0x10^5
Freq.error = 0.18
F.
pN2-6 + MeNC Activation (short freezing intervals)
n = 4.2x10^5
Freq.error = 0.17
G.
pN2-6 + MeNC Activation (short freezing activation intervals and 4°C solution-phase activation intervals)
n = 5.3x10^5
Freq.error = 0.25
H.
pN2-6 + MeNC Activation (short freezing and short solution-phase activation intervals)
n = 3.7x10^5
Freq.error = 0.18
I.
pN2-6 + MeNC Activation (short freezing activation intervals and short solution-phase activation intervals at 4°C)
n = 4.1x10^5
Freq.error = 0.27
0 0.06 0.12 0.18 0.24 0.3
1 2 3 4 5 6 7 8 9 10

**Supplementary Figure 12. Conditions that increase bridged mononucleotide-oligonucleotide formation or lack the full oligo mix increase the frequency of mismatches. A-I.** The cumulative sum frequency distributions of correct and incorrect nucleotide incorporations normalized to each position for experiments with 20.5 mM 2AI, 20 mM rN, the indicated oligonucleotide mixture, eutectic MeNC activation, and solution-phase MeNC activation (see Supplementary Table 2, entries S12A-I, for all exact conditions). **A**. Reducing the interval at which MeNC is added increases errors. **B.** Changing the distribution of oligonucleotides from 1 mM total with progressively lower concentrations of longer oligonucleotides (as in the oligo mix) to a flat distribution of 16 µM total with 1 µM each oligonucleotide increases errors. **C.** Using only oligos $pN_{2-6}$ is also error-prone. **D.** Using only oligos $pN_{7-16}$ is also error-prone. **E.** Reducing the temperature of the solution-phase reaction from 23°C (standard) to 4°C reduces yields slightly but has no effect on errors (compare to (C)). **F-I.** Various combinations of the conditions tested above, as indicated, using $pN_{2-6}$, all result in high error frequencies.

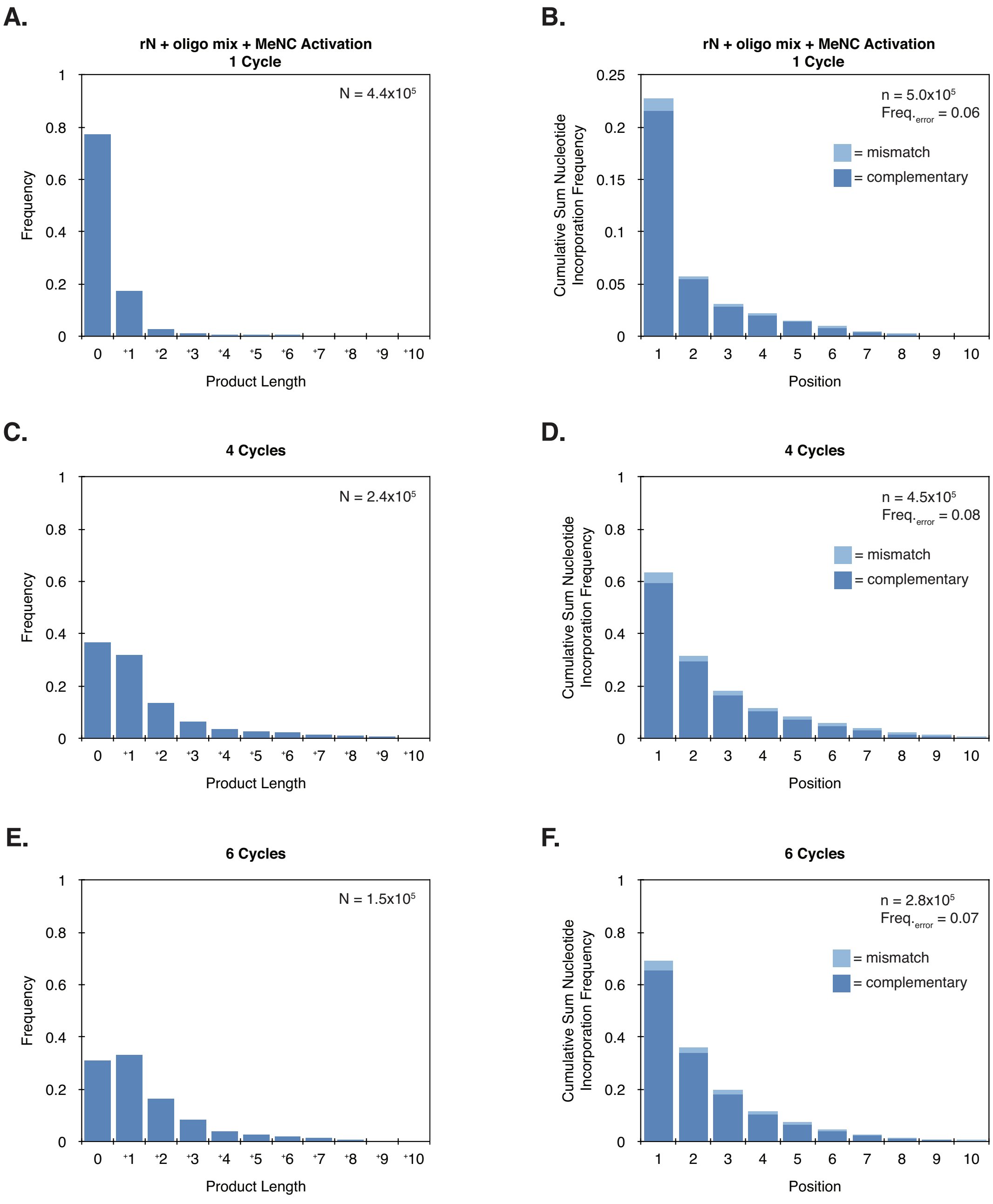

**Supplementary Figure 13. Cycles of eutectic and solution-phase activation progressively increase yields. A-F.** Product length distributions and cumulative sum frequency distributions of correct and incorrect nucleotide incorporations of primer extension with 20 mM rN + 1 mM oligo mix ($pN_{2-16}$) + *in situ* MeNC activation **A-B.** cycled once (same data as Supplementary Figure 3E-F, for comparison), **C-D.** four times, or **E-F.** six times. Note the change in the y-axis scale for the cumulative sum histograms between one and four cycles. By four cycles, more than half of the sequencing hairpins have been extended. **F.** The frequency of incorrect incorporations is 0.069 after six cycles, compared to 0.063 for one cycle (B).

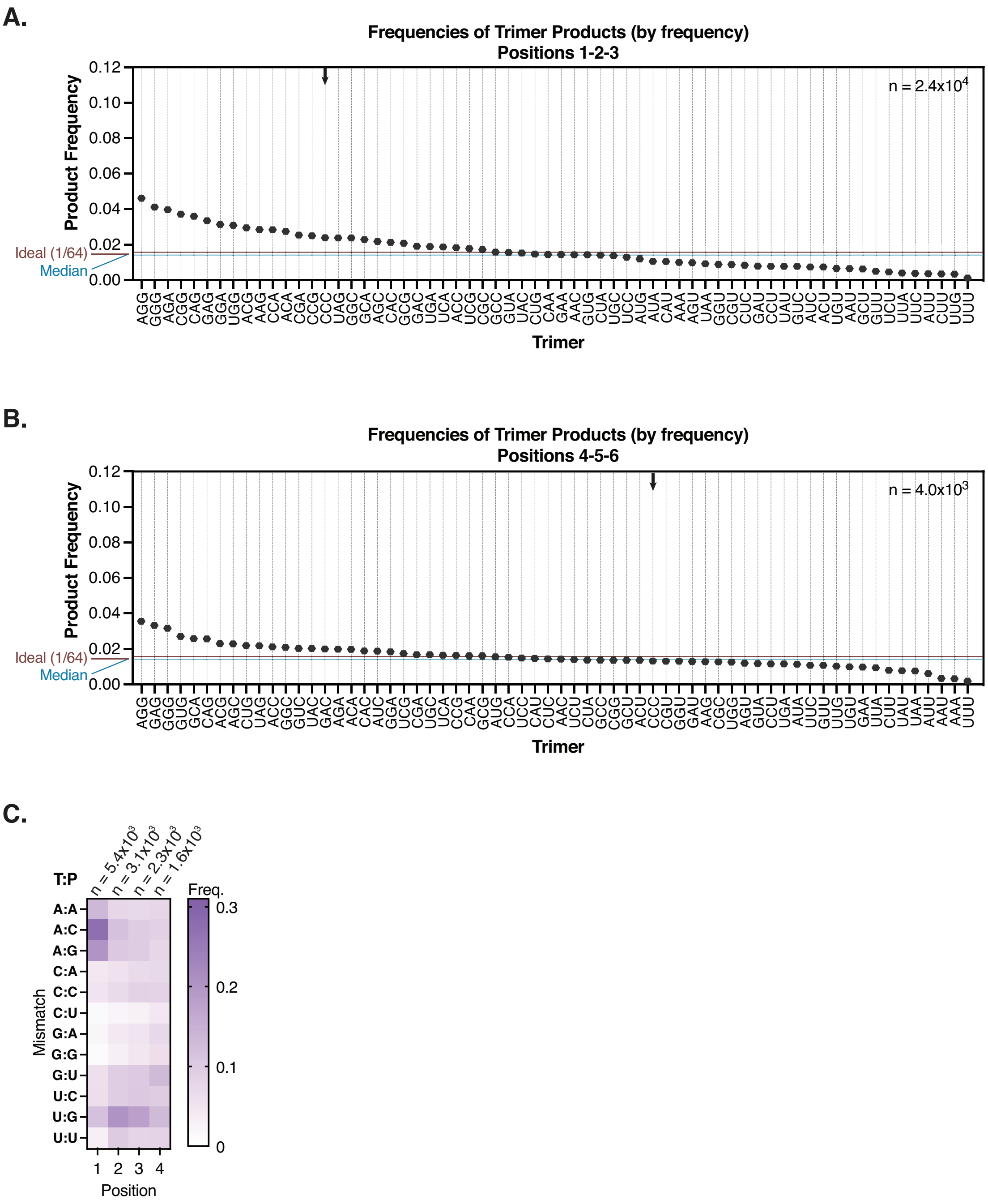

**Supplementary Figure 14. Additional product features for six cycles of primer extension. A-C.** Product sequence features of primer extension with 20 mM rN + 1 mM oligo mix (pN$_{2\text{-}16}$) + *in situ* MeNC activation cycled six times. **A.** The quantification of trimer frequencies for positions 1-2-3, ordered by trimer frequency (same data as in Figure 5E). The median frequency is 0.014. The ideal frequency for each trimer is 1/64 = 0.016. **B.** The quantification of trimer frequencies for positions 4-5-6, ordered by trimer frequency (same data as in Figure 5F). The median frequency is 0.014. **C.** The distribution of mismatches (listed as **T**emplate:**P**roduct) shows features associated with both bridged dinucleotide primer extension and ligation (see Supplementary Figure 4B).

### EXTENDED MATERIAL AND METHODS

**Synthesis of 2-aminoimidazole-activated monoribonucleotides** (1)

0.63 mmole (1 equivalent) of the nucleoside-5′-monophosphate free acid (Santa Cruz Biotechnology or ACROS Organics) and 5 equivalents of 2-aminoimidazole hydrochloride (Combi-Blocks) were dissolved in water, the pH adjusted to 5.5 with NaOH, and the mixture lyophilized. 30 ml dry dimethyl sulfoxide (DMSO, Sigma-Aldrich) and 1.2 ml dry triethylamine (TEA, Sigma-Aldrich) were mixed under argon. To this was added the lyophilized nucleoside-5′-monophosphate and 2-aminoimidazole hydrochloride, 9 equivalents of triphenylphosphine (TPP, Sigma-Aldrich), and 10 equivalents of 2,2′-dipyridyldisulfide (DPDS, Combi-Blocks). After 30 minutes the mixture was poured into an ice-chilled solution of 400 ml acetone (Fisher Scientific), 250 ml diethyl ether (Fisher Scientific), 30 ml TEA, and 1.6 ml acetone saturated with $NaClO_4$ (Sigma-Aldrich). After 30 minutes the supernatant was removed. The remaining mixture with the flocculant precipitate was centrifuged, the supernatant discarded, and the pellet washed in a solution of 1:8.3:13.3 of TEA:diethyl ether:acetone and centrifuged again. The wash was repeated twice with just acetone, and the pellets dried overnight under vacuum. Each sample was dissolved in 5 ml water, and the combined volume purified by reverse-phase flash chromatography (CombiFlash, Teledyne ISCO) over a 50 g RediSep Rf Gold C18Aq column (Teledyne ISCO) using gradient elution between (A) water and (B) acetonitrile. Target fractions were placed on ice, pooled, adjusted to pH 10 with NaOH, lyophilized, and stored at -80°C. Purified material was resuspended in water, the nucleotide concentration measured by a NanoDrop 2000c spectrophotometer (ThermoFisher Scientific), and stock aliquots were prepared with equimolar concentrations of each 2AI-activated nucleotide (2AIrA + 2AIrU + 2AIrG + 2AIrC = 2AIrN). Aliquots were lyophilized, stored at -80°C, and resuspended in water immediately prior to use.

**Working stocks of unactivated nucleotides**

A nominally 800 mM equimolar mixture of rA, rU, rG, and rC nucleoside-5′-monophosphate $Na^+$ salts (rN, Sigma-Aldrich or ACROS Organics) was prepared as a stock in water and adjusted to pH 8. To ensure that direct comparisons can be made with experiments that used the stock mixture of all four pre-activated nucleotides (2AIrN, as prepared above), the concentration value of the rN mix was adjusted to agree with the concentration of 2AIrN as determined by $A_{260}$ on a NanoDrop 2000c spectrophotometer.

**Oligonucleotides**

**General** (3)

Oligonucleotides synthesised in-house were prepared on an ABI Expedite instrument (standard phosphoramidites from ChemGenes; special phosphoramidites, synthesis reagents and supports from Glen Research) at the 1 µmole scale unless otherwise indicated with DMT-on for purification. Oligo stocks were stored in 200 mM HEPES pH 7.5 at -20 or -80°C.

**Unphosphorylated oligonucleotides** (3)

The product was cleaved from the support with 1.3 ml of a 1:1 mixture of 28-30% ammonium hydroxide and 40% aqueous methylamine (AMA). The bases were deprotected by incubation at 65°C for 10 minutes, then vacuum centrifuge concentrated for two hours until ~50% of the initial volume was left. The sample was flash frozen and lyophilized overnight at -20°C. The lyophilized material was resuspended in 115 µl DMSO, 60 µl TEA, and 75 µl TEA/3HF, incubated at 65°C for 2.5 hours, and allowed to come to room temperature. The oligo was then purified with a GlenPak column (Glen Research), eluted with 200 mM ammonium bicarbonate (Sigma-Aldrich, from a syringe-filtered 1 M stock) + 30% v/v acetonitrile (Fisher Scientific) (the first one or two drops and the final foam were discarded), vacuum centrifuge concentrated for 45 minutes, flash frozen in liquid nitrogen, and lyophilized overnight at -20°C.

The lyophilized material was resuspended in 500 µl TE, pH 7 and PAGE-purified. 50 µl of the oligo stock was mixed with 50 µl Urea Load Buffer (8.3 M urea [Sigma-Aldrich], 1.3x TBE buffer [from a 10x autoclaved stock], 75 µM bromophenol blue [Sigma-Aldrich, from a 7.5 mM stock in DMSO], 880 µM orange G [Sigma-Aldrich, from an 88 mM stock in DMSO], syringe-filtered) and subjected to PAGE at 10 W for 15 minutes, then 25 W for 1 hour. One of the two glass plates was removed (leaving the other for the gel to rest on), and 10 ml TBE + 1 µl SYBR Gold Nucleic Acid Gel Stain (Invitrogen) was poured on top of the gel and allowed to incubate for 3-4 minutes before being rinsed off with Milli-Q water. The gel was imaged with a blue light transilluminator (Safe Imager 2.0, ThermoFisher Scientific): the target material typically appeared as unstained due to its extremely high concentration relative to well-stained higher (not fully deprotected) and lower (truncated) molecular weight side-products. The target band was excised with a clean razor, purified with a ZR small-RNA PAGE Recovery Kit (Zymo Research), eluted in 30 µl TE, pH 7 and quantified with a NanoDrop spectrophotometer. Product identity was confirmed by Time-Of-Flight Liquid Chromatography-coupled Mass Spectroscopy (TOF LC-MS, Agilent 6230), except in the case of hairpin constructs, which were too large for accurate mass spectroscopy analysis.

**5′ phosphorylated oligonucleotides (except $pN_2$)**

We sought to avoid potential biases in the phosphorylation of random-sequence oligonucleotides by chemical phosphorylating agents and developed a new approach to easily synthesize and prepare 5′ phosphorylated oligonucleotides. Reverse amidites (these synthesize in the 5′ ⟶ 3′ direction) were purchased from ChemGenes; for random-sequence oligonucleotide syntheses, they were mixed in a 1:1:1:1 ratio. Synthesis was performed on a Glen Research chemical phosphorylating agent (CPA) support (so-called 3′-phosphorylating support, because this manufacturer only supplies regular amidites that synthesize in the 3′ ⟶ 5′ direction). This synthesis results in the 5′ end of the newly synthesized oligo attached to the CPA support so that conversion to the phosphate can occur during AMA cleavage while leaving the 3′ end free for a terminal DMT-on, to be used as a handle for later purification. Downstream work up is the same as for unphosphorylated oligos except incubation in AMA after elution is for 30 minutes instead of 10 minutes, and 2% TFA incubation is for 2 x 5 minutes. (3′ DMT is more stable than 5′ DMT and requires a longer acid incubation for cleavage. This does not lead to depurination.) $pN_3$ and $pN_4$ were prepared at the 15 µmole scale, and the products cleaved from the support with 5 ml AMA instead of 1.3 ml. The remainder of the work up was as above, but split four ways to account for the greater quantity of product. Random-sequence oligonucleotides could not be gel purified because those used in this study were too short for gel visualization, and they were required in high concentrations. Random-sequence oligonucleotides were eluted from the GlenPak column in 5 mM TEAB and 20% acetonitrile.

This protocol was validated by LC-MS for two short defined-sequence oligos. pCGG showed one primary UV absorbance peak: m/z expected = 1012.137, measured = 1012.14. pGGU: showed one primary UV absorbance peak: m/z expected = 1013.121; measured = 1013.12. A very minor peak was found to correspond to pGU (m/z = 668.07).

**pN$_2$**

5′ phosphorylated dimers are too short to retain on a GlenPak column after TFA cleavage and are washed out, and lost, by the water quench of the acid. Instead, pN$_2$ was prepared as the other random-sequence oligos and loaded on a GlenPack column to clean up the HF reaction (the DMT group is sufficient to retain it, also ensuring that failure sequences without a DMT group are washed away). However, the on-column 2% TFA cleavage step was skipped, and the otherwise washed and deprotected but still DMT-on dimers were eluted in 5 mM TEAB and 40% acetonitrile. The eluate was vacuum centrifuge concentrated for 45 minutes, flash frozen in liquid nitrogen and lyophilized overnight at -20°C. The lyophilized powder was resuspended in 100 µl DMSO and 300 µl 2% TFA, incubated for 15 minutes, quenched with 300 µl 1 M TEAB, and vacuum centrifuge concentrated for 30 minutes. The concentrated mixture was purified by reverse-phase flash chromatography (CombiFlash, Teledyne ISCO) over a 5.5 g RediSep Rf Gold C18Aq column (Teledyne ISCO) using gradient elution between (A) 5 mM TEAB and (B) acetonitrile. Peaks were monitored by 260 nm and 230 nm absorbance, with the target RNA peak showing high 260 nm absorbance and organic contaminants showing high 230 nm absorbance. The target fraction eluted at ~10% acetonitrile, and was vacuum centrifuge concentrated for ~30 minutes and lyophilized.

### Supplementary Table 2, Oligonucleotides

| Oligo Name | Type | Source | Sequence (5′-3′; all termini free -OH unless otherwise noted) | Notes |
|---|---|---|---|---|
| 6N | RNA/DNA | In-house† | GUUCAGAGUUCUACAGUCCGACGAUCdT(-NPOM)CdT(-NPOM)ANNNNNNGCAUGCGACUAAACGUCGCAUGC | Sequencing hairpin with a random 6-base template (1,3) |
| 18N | RNA/DNA | In-house | GUUCAGAGUUCUACAGUCCGACGAUCdT(-NPOM)CdT(-NPOM)ANNNNNNNNNNNNNNNNNNGCAUGCGACUAAACGUCGCAUGC | Sequencing hairpin with a random 18-base template |
| CTB | RNA/DNA | In-house | GUUCAGAGUUCUACAGUCCGACGAUCdT(-NPOM)CdT(-NPOM)AAAACCGGCAUGCGACUAAACGUCGCAUGC | Sequencing hairpin with "GCCAAA" template (3) |
| 5′ Handle Block | RNA | IDT⊥ | GUCGGACUGUAGAACUCUGAA-dideoxyC | Binds 5′ end of sequencing hairpin and prevents it from interfering with the template sequence (3) |
| $N_2$ - $N_{16}$ | RNA | In-house | $pN_{(2\text{-}16)}$ | 5′-phosphorylated random-sequence oligonucleotides, of lengths 2 through 16. Each length was prepared separately. |
| pCGG | RNA | In-house | pCGG | For use with CTB. See also Extended Materials and Methods |
| $N_6$ 5′-G | RNA | In-house | p(A,U,C)NNNNN | 5′-phosphorylated random-sequence hexamer except 5′-terminal position lacks rG |
| RT Handle | DNA | IDT | App-AGATCGGAAGAGCACACGTCT-dideoxyC | Template for the R(everse) T(ranscription) Primer |
| RT Primer | DNA | IDT | AGACGTGTGCTCTTCCGATCT | |
| PCR Primer 1 (NEBNext SR Primer for Illumina) | DNA | NEB | AATGATACGGCGACCACCGAGATCTACACGTTCAGAGTTCTACAGTCCG-s-A | |
| PCR Primer 2 (NEBNext Index | DNA | NEB | CAAGCAGAAGACGGCATACGAGAT(6-base index)GTGACTGGAGTTCAGACGTGTGCTCTTCCGATC-s-T | |

| |
|---|
| Primer for Illumina) |

† See Material and Methods and (3) for details on in-house oligo synthesis and purification
dT(-NPOM) = NPOM-caged deoxyT (2)
N = rA, rU, rG, or rC
⊥ IDT oligos ordered as RNase-free HPLC-purified
App = riboA 5′-adenylation
-s- = thiol backbone linkage to inhibit exonucleases

### Supplementary Table 3, Sequencing Experiments

| File Designation | Description | Conditions | Notes | Figure(s) |
|---|---|---|---|---|
| **U_6N130Mg** | Sequencing hairpin template + pre-activated nucleotides exposed to 23°C MeNC activation for 24 hours | 100 mM HEPES-Na pH 8, 1 µM 6N sequencing hairpin, 1.2 µM 5′ Handle Block oligo, 20 mM rNMP-2AI (5 mM each of rAMP-2AI, rUMP-2AI, rGMP-2AI, and rCMP-2AI), 30 mM $MgCl_2$, and 260 mM 2MBA + 200 mM MeNC incubated for 24 hours at 23°C | Under these conditions the activation chemistry drives bridged dinucleotide formation. Serves as a standard for comparison; uses previously-established buffer conditions for MeNC-based activation (5,6). | 2, S3, S4 |
| **Y_18NC4** | Sequencing hairpin template + reaction buffer, MeNC activation chemistry, and no reactants | 100 mM HEPES-Na pH 8, 1 µM 18N sequencing hairpin, 1.2 µM 5′ Handle Block oligo, 30 mM $MgCl_2$, and 260 mM 2MBA + 200 mM MeNC incubated for 24 hours at 23°C | Control template for the data analysis software (3) for experiments with 18N sequencing hairpin plus MeNC activation chemistry. Note that MeNC activation is not expected to affect the RNA under these conditions and this control has been included for completeness. | |
| **CCN_18N1a** | Sequencing hairpin template, mixture of initially unactivated random-sequence oligonucleotides of increasing lengths ($pN_{2\text{-}16}$) in decreasing concentrations, and initially unactivated nucleotides exposed to four rounds of eutectic MeNC activation followed by four rounds of 23°C MeNC activation at 24 hour intervals | 50 mM HEPES-Na pH 8, 1 µM 18N sequencing hairpin, 1.2 µM 5′ Handle Block oligo, 1 mM total oligonucleotide mixture (oligo mix) (500 µM $N_2$, 250 µM $N_3$, 125 µM $N_4$, 62.5 µM $N_5$, 31.3 µM $N_6$, 15.6 µM $N_7$, 7.81 µM $N_8$, 3.91 µM $N_9$, 1.95 µM $N_{10}$, 0.98 µM $N_{11}$, 0.49 µM $N_{12}$, 0.24 µM $N_{13}$, 0.12 µM $N_{14}$, 0.06 µM $N_{15}$, and 0.03 µM $N_{16}$), 20 mM rNMP (5 mM each of rAMP, rUMP, rGMP, and rCMP), 20.5 mM 2-aminoimidazole (2AI), and 30 mM $MgCl_2$; 260 mM total 2-methylbutyraldehyde (2MBA) added once + | The oligo mix was prepared in advance as a 5 mM stock solution. | 2, S3, S4, S13 |

| | | | | |
|---|---|---|---|---|
| | | 50 mM methyl isocyanide (MeNC) added at each brief thaw between four frozen incubations (4 x 24 hours at -17°C); thaw + 24 hours at 23°C; 260 mM total 2MBA added once + 50 mM MeNC added four times (4 x 24 hours at 23°C) | | |
| **CCN_18NN3** | Sequencing hairpin template, initially unactivated $pN_3$, and initially unactivated nucleotides exposed to four rounds of eutectic MeNC activation followed by four rounds of 23°C MeNC activation at 24 hour intervals | 50 mM HEPES-Na pH 8, 1 µM 18N sequencing hairpin, 1.2 µM 5′ Handle Block oligo, 250 µM $N_3$, 20 mM rNMP (5 mM each of rAMP, rUMP, rGMP, and rCMP), 20.5 mM 2-aminoimidazole (2AI), and 30 mM $MgCl_2$; 260 mM total 2-methylbutyraldehyde (2MBA) added once + 50 mM methyl isocyanide (MeNC) added at each brief thaw between four frozen incubations (4 x 24 hours at -17°C); thaw + 24 hours at 23°C; 260 mM total 2MBA added once + 50 mM MeNC added four times (4 x 24 hours at 23°C) | | S3 |
| **Q_6N(5, 10, 20, or 40)un & S_6N(80 or 160)un** | Pre-activated nucleotides + increasing concentrations of unactivated nucleotide competitor | 200 mM HEPES-Na pH 8, 1 µM 6N sequencing hairpin, 1.2 µM 5′ Handle Block oligo, 20 mM rNMP-2AI (5 mM each of rAMP-2AI, rUMP-2AI, rGMP-2AI, and rCMP-2AI), 3.6, 7.2, 14.4, 28.8, 57.6, or 115.2 mM unactivated rNMP competitor (named 6N5un, 6N10un, 6N20un, 6N40un, 6N80un, and 6N160un respectively), and 50 mM $MgCl_2$ incubated for 24 hours at 23°C | See Materials and Methods (1) for concentrations of unactivated rNMPs | S5 |
| **W_18NC2** | Sequencing hairpin template + reaction buffer and no reactants | 100 mM HEPES-Na pH 8, 1 µM 18N sequencing hairpin, 1.2 µM 5′ Handle Block oligo, and 30 mM $MgCl_2$ incubated for 24 hours at 23°C | Control template for the data analysis software (3) for experiments with 18N sequencing hairpin and no MeNC activation chemistry | |
| **Z_18NOlig2, W_18NOlig(3, 4, ..., or 14) & Z_18NOlig(15 or 16)** | Pre-activated nucleotides + unactivated random-sequence oligonucleotide competitors of increasing lengths ($pN_{2-16}$) in | 100 mM HEPES-Na pH 8, 1 µM 18N sequencing hairpin, 1.2 µM 5′ Handle Block oligo, 20 mM rNMP-2AI (5 mM each of rAMP-2AI, rUMP-2AI, rGMP-2AI, and rCMP-2AI), unactivated 5′-phosphorylated random- | | S5, S6 |

| | | | | |
|---|---|---|---|---|
| | decreasing concentrations, one at a time | sequence oligonucleotide competitor, one length oligonucleotide at a time (500 µM $pN_2$; 250 µM $pN_3$; 125 µM $pN_4$; 62.5 µM $pN_5$; 31.3 µM $pN_6$; 15.6 µM $pN_7$; 7.8 µM $pN_8$; 3.9 µM $pN_9$; 2 µM $pN_{10}$; 0.98 µM $pN_{11}$; 0.49 µM $pN_{12}$; 0.24 µM $pN_{13}$; 0.12 µM $pN_{14}$; 0.06 µM $pN_{15}$; or 0.03 µM $pN_{16}$), and 30 mM $MgCl_2$ incubated for 24 hours at 23°C | | |
| **Z_18NOlig6-9-12-15** | Pre-activated nucleotides + mixture of $pN_6$, $pN_9$, $pN_{12}$, and $pN_{15}$ | 100 mM HEPES-Na pH 8, 1 µM 18N sequencing hairpin, 1.2 µM 5′ Handle Block oligo, 20 mM rNMP-2AI (5 mM each of rAMP-2AI, rUMP-2AI, rGMP-2AI, and rCMP-2AI), 35.8 µM mixture of unactivated 5′-phosphorylated random-sequence oligonucleotides (31.3 µM $pN_6$, 3.9 µM $pN_9$, 0.49 µM $pN_{12}$, and 0.06 µM $pN_{15}$), and 30 mM $MgCl_2$ incubated for 24 hours at 23°C | The different length oligonucleotides were mixed in advance. | S5, S7 |
| **Z_18NOlig6hlfX** | Pre-activated nucleotides + $pN_6$ at half the standard concentration | 100 mM HEPES-Na pH 8, 1 µM 18N sequencing hairpin, 1.2 µM 5′ Handle Block oligo, 20 mM rNMP-2AI (5 mM each of rAMP-2AI, rUMP-2AI, rGMP-2AI, and rCMP-2AI), 15.7 µM $pN_6$, and 30 mM $MgCl_2$ incubated for 24 hours at 23°C | | S6 |
| **Z_18NOlig6NOrG** | Pre-activated rA, rU, and rC (no rG) + $pN_6$ | 100 mM HEPES-Na pH 8, 1 µM 18N sequencing hairpin, 1.2 µM 5′ Handle Block oligo, 20 mM rNMP-2AI (6.67 mM each of rAMP-2AI, rUMP-2AI, and rCMP-2AI), 31.3 µM $pN_6$, and 30 mM $MgCl_2$ incubated for 24 hours at 23°C | | S6 |
| **Z_18NOlig65primenoG** | Pre-activated nucleotides + unactivated random-sequence hexamer competitor lacking rG at 5′ terminal positon | 100 mM HEPES-Na pH 8, 1 µM 18N sequencing hairpin, 1.2 µM 5′ Handle Block oligo, 20 mM rNMP-2AI (5 mM each of rAMP-2AI, rUMP-2AI, rGMP-2AI, and rCMP-2AI), 31.3 µM $N_6$ 5′-G, and 30 mM $MgCl_2$ incubated for 24 hours at 23°C | | S6 |
| **Z_18NOligMix** | Pre-activated nucleotides + mixture of unactivated random-sequence oligonucleotides of | 100 mM HEPES-Na pH 8, 1 µM 18N sequencing hairpin, 1.2 µM 5′ Handle Block oligo, 20 mM rNMP-2AI (5 mM each of rAMP- | The oligo mix was prepared in advance as a 5 mM stock solution. | S5, 3 |

| | | | | |
|---|---|---|---|---|
| | increasing lengths ($pN_{2-16}$) in decreasing concentrations | 2AI, rUMP-2AI, rGMP-2AI, and rCMP-2AI), 1 mM mixture of unactivated 5′-phosphorylated random-sequence oligonucleotides (oligo mix) (500 µM $pN_2$, 250 µM $pN_3$, 125 µM $pN_4$, 62.5 µM $pN_5$, 31.3 µM $pN_6$, 15.6 µM $pN_7$, 7.8 µM $pN_8$, 3.9 µM $pN_9$, 2 µM $pN_{10}$, 0.98 µM $pN_{11}$, 0.49 µM $pN_{12}$, 0.24 µM $pN_{13}$, 0.12 µM $pN_{14}$, 0.06 µM $pN_{15}$, and 0.03 µM $pN_{16}$), and 30 mM $MgCl_2$ incubated for 24 hours at 23°C | | |
| **Z_18NOligMixtwoX** | Pre-activated nucleotides + mixture of unactivated random-sequence oligonucleotides of increasing lengths ($pN_{2-16}$) in decreasing concentrations at twice the standard total concentration | 100 mM HEPES-Na pH 8, 1 µM 18N sequencing hairpin, 1.2 µM 5′ Handle Block oligo, 20 mM rNMP-2AI (5 mM each of rAMP-2AI, rUMP-2AI, rGMP-2AI, and rCMP-2AI), 2 mM mixture of unactivated 5′-phosphorylated random-sequence oligonucleotides (oligo mix) (1 mM $pN_2$, 500 µM $pN_3$, 250 µM $pN_4$, 125 µM $pN_5$, 62.5 µM $pN_6$, 31.3 µM $pN_7$, 15.6 µM $pN_8$, 7.8 µM $pN_9$, 3.9 µM $pN_{10}$, 2 µM $pN_{11}$, 0.98 µM $pN_{12}$, 0.49 µM $pN_{13}$, 0.24 µM $pN_{14}$, 0.12 µM $pN_{15}$, and 0.06 µM $pN_{16}$), and 30 mM $MgCl_2$ incubated for 24 hours at 23°C | The oligo mix was prepared in advance as a 5 mM stock solution. | S5, S7 |
| **Z_18NOligMixhlfX** | Pre-activated nucleotides + mixture of unactivated random-sequence oligonucleotides of increasing lengths ($pN_{2-16}$) in decreasing concentrations at half the standard total concentration | 100 mM HEPES-Na pH 8, 1 µM 18N sequencing hairpin, 1.2 µM 5′ Handle Block oligo, 20 mM rNMP-2AI (5 mM each of rAMP-2AI, rUMP-2AI, rGMP-2AI, and rCMP-2AI), 0.5 mM mixture of unactivated 5′-phosphorylated random-sequence oligonucleotides (oligo mix) (250 µM $pN_2$, 125 µM $pN_3$, 62.5 µM $pN_4$, 31.3 µM $pN_5$, 15.6 µM $pN_6$, 7.8 µM $pN_7$, 3.9 µM $pN_8$, 2 µM $pN_9$, 0.98 µM $pN_{10}$, 0.49 µM $pN_{11}$, 0.24 µM $pN_{12}$, 0.12 µM $pN_{13}$, 0.06 µM $pN_{14}$, 0.03 µM $pN_{15}$, and 0.015 µM $pN_{16}$), and 30 mM $MgCl_2$ incubated for 24 hours at 23°C | The oligo mix was prepared in advance as a 5 mM stock solution. | S5, S7 |
| **CCN_18NOligMixNmix** | Sequencing hairpin template, mixture of initially unactivated random-sequence oligonucleotides of increasing | 50 mM HEPES-Na pH 8, 1 µM 18N sequencing hairpin, 1.2 µM 5′ Handle Block oligo, 1 mM total oligonucleotide mixture (oligo mix) (500 µM $N_2$, 250 µM $N_3$, 125 µM | The oligo mix was prepared in advance as a 5 mM stock solution. | 3 |

| | | | | |
|---|---|---|---|---|
| | lengths ($pN_{2-16}$) in decreasing concentrations, and pre-activated nucleotides exposed to 23°C MeNC activation for 24 hours | $N_4$, 62.5 µM $N_5$, 31.3 µM $N_6$, 15.6 µM $N_7$, 7.81 µM $N_8$, 3.91 µM $N_9$, 1.95 µM $N_{10}$, 0.98 µM $N_{11}$, 0.49 µM $N_{12}$, 0.24 µM $N_{13}$, 0.12 µM $N_{14}$, 0.06 µM $N_{15}$, and 0.03 µM $N_{16}$), 20 mM rNMP-2AI (5 mM each of rAMP-2AI, rUMP-2AI, rGMP-2AI, and rCMP-2AI), 30 mM $MgCl_2$, 260 mM 2-methylbutyraldehyde (2MBA), and 200 mM methyl isocyanide (MeNC) incubated for 24 hours at 23°C | | |
| **DDA_18N1a** | Sequencing hairpin template, mixture of initially unactivated random-sequence oligonucleotides of increasing lengths ($pN_{2-16}$) in decreasing concentrations, and initially unactivated nucleotides exposed to four rounds of eutectic high concentration MeNC activation followed by four rounds of 23°C high concentration MeNC activation at 24 hour intervals | 50 mM HEPES-Na pH 8, 1 µM 18N sequencing hairpin, 1.2 µM 5′ Handle Block oligo, 1 mM total oligonucleotide mixture (oligo mix) (500 µM $N_2$, 250 µM $N_3$, 125 µM $N_4$, 62.5 µM $N_5$, 31.3 µM $N_6$, 15.6 µM $N_7$, 7.81 µM $N_8$, 3.91 µM $N_9$, 1.95 µM $N_{10}$, 0.98 µM $N_{11}$, 0.49 µM $N_{12}$, 0.24 µM $N_{13}$, 0.12 µM $N_{14}$, 0.06 µM $N_{15}$, and 0.03 µM $N_{16}$), 20 mM rNMP (5 mM each of rAMP, rUMP, rGMP, and rCMP), 20.5 mM 2-aminoimidazole (2AI), and 30 mM $MgCl_2$; 260 mM total 2-methylbutyraldehyde (2MBA) added once + 100 mM methyl isocyanide (MeNC) added at each brief thaw between four frozen incubations (4 x 24 hours at -17°C); thaw + 24 hours at 23°C; 260 mM total 2MBA added once + 100 mM MeNC added four times (4 x 24 hours at 23°C) | The oligo mix was prepared in advance as a 5 mM stock solution. Concentration of MeNC 2x that used in CCN_18N1a | S8 |
| **DDA_18N(2, 3, ..., or 8) & DDB_18N(9, 10, ..., or 16)** | Sequencing hairpin template, initially unactivated random-sequence oligonucleotides of increasing lengths ($pN_{2-16}$) in decreasing concentrations, one at a time, and initially unactivated nucleotides exposed to four rounds of eutectic high concentration MeNC activation followed by four rounds of 23°C | 50 mM HEPES-Na pH 8, 1 µM 18N sequencing hairpin, 1.2 µM 5′ Handle Block oligo, 5′-phosphorylated random-sequence oligonucleotide one length oligonucleotide at a time (500 µM $pN_2$; 250 µM $pN_3$; 125 µM $pN_4$; 62.5 µM $pN_5$; 31.3 µM $pN_6$; 15.6 µM $pN_7$; 7.8 µM $pN_8$; 3.9 µM $pN_9$; 2 µM $pN_{10}$; 0.98 µM $pN_{11}$; 0.49 µM $pN_{12}$; 0.24 µM $pN_{13}$; 0.12 µM $pN_{14}$; 0.06 µM $pN_{15}$; or 0.03 µM $pN_{16}$), 20 mM rNMP (5 mM each of rAMP, rUMP, rGMP, and rCMP), 20.5 mM 2- | Concentration of MeNC 2x that used in CCN_18N1a | S8 |

| | | | | |
|---|---|---|---|---|
| | high concentration MeNC activation at 24 hour intervals | aminoimidazole (2AI), and 30 mM $MgCl_2$; 260 mM total 2-methylbutyraldehyde (2MBA) added once + 100 mM methyl isocyanide (MeNC) added at each brief thaw between four frozen incubations (4 x 24 hours at -17°C); thaw + 24 hours at 23°C; 260 mM total 2MBA added once + 100 mM MeNC added four times (4 x 24 hours at 23°C) | | |
| **AA_CTBLig(50, 100, or 200)** | Defined-template (GCCAAA) sequencing hairpin + pCGG exposed to 23°C MeNC activation for 24 hours with 50, 100, or 200 mM 2AI | 100 mM HEPES-Na pH 8, 1 µM CTB sequencing hairpin, 1.2 µM 5′ Handle Block oligo, 250 µM pCGG, 50, 100, or 200 mM 2-aminoimidazole (2AI), 30 mM $MgCl_2$, 260 mM 2-methylbutyraldehyde (2MBA), and 200 mM methyl isocyanide (MeNC) incubated for 24 hours at 23°C | Ligation-only conditions with 2-aminomidazole as the activating group | ST1 |
| **Y_CTBLigB** | Defined-template (GCCAAA) sequencing hairpin + pCGG exposed to four rounds of eutectic MeNC activation at 24 hour intervals followed by a 24 hour incubation at 23°C | 100 mM HEPES-Na pH 8, 1 µM CTB sequencing hairpin, 1.2 µM 5′ Handle Block oligo, 250 µM pCGG, 20 mM 2-aminoimidazole (2AI), and 30 mM $MgCl_2$; 260 mM total 2-methylbutyraldehyde (2MBA) added once + 50 mM methyl isocyanide (MeNC) added at each brief thaw between four frozen incubations (4 x 24 hours at -17°C); thaw + 24 hours at 23°C | Ligation-only conditions with 2-aminoimidazole as the activating group | ST1 |
| **Y_18NOlig3L** | Sequencing hairpin tempate + $pN_3$ exposed to four rounds of eutectic MeNC activation at 24 hour intervals followed by a 24 hour incubation at 23°C | 100 mM HEPES-Na pH 8, 1 µM 18N sequencing hairpin, 1.2 µM 5′ Handle Block oligo, 250 µM $pN_3$, 20 mM 2-aminoimidazole (2AI), and 30 mM $MgCl_2$; 260 mM total 2-methylbutyraldehyde (2MBA) added once + 50 mM methyl isocyanide (MeNC) added at each brief thaw between four frozen incubations (4 x 24 hours at -17°C); thaw + 24 hours at 23°C | Ligation-only conditions with 2-aminoimidazole as the activating group | ST1 |
| **CCN_18NLigN2** | Sequencing hairpin template + pN2 exposed to 23°C MeNC activation for 24 hours with 200 mM 2MI | 50 mM HEPES-Na pH 8, 1 µM 18N sequencing hairpin, 1.2 µM 5′ Handle Block oligo, 500 µM $pN_2$, 200 mM 2-methylimidazole (2MI), 50 mM $MgCl_2$, 260 mM 2-methylbutyraldehyde (2MBA), and 200 | Ligation-only conditions with 2-methylimidazole as the activating group | ST1 |

| | | | | |
|---|---|---|---|---|
| | | mM methyl isocyanide (MeNC) incubated for 24 hours at 23°C | | |
| **CCN_18NLigN3** | Sequencing hairpin template + $pN_3$ exposed to 23°C MeNC activation for 24 hours with 200 mM 2MI | 50 mM HEPES-Na pH 8, 1 µM 18N sequencing hairpin, 1.2 µM 5′ Handle Block oligo, 250 µM $pN_3$, 200 mM 2-methylimidazole (2MI), 50 mM $MgCl_2$, 260 mM 2-methylbutyraldehyde (2MBA), and 200 mM methyl isocyanide (MeNC) incubated for 24 hours at 23°C | Ligation-only conditions with 2-methylimidazole as the activating group | ST1 |
| **EEA_18NL1a** | Sequencing hairpin template and mixture of initially unactivated random-sequence oligonucleotides of increasing lengths ($pN_{2-16}$) in decreasing concentrations exposed to 23°C high concentration MeNC activation for 48 hours with 200 mM 1MI | 100 mM HEPES-Na pH 8, 1 µM 18N sequencing hairpin, 1.2 µM 5′ Handle Block oligo, 1 mM total oligonucleotide mixture (oligo mix) (500 µM $N_2$, 250 µM $N_3$, 125 µM $N_4$, 62.5 µM $N_5$, 31.3 µM $N_6$, 15.6 µM $N_7$, 7.81 µM $N_8$, 3.91 µM $N_9$, 1.95 µM $N_{10}$, 0.98 µM $N_{11}$, 0.49 µM $N_{12}$, 0.24 µM $N_{13}$, 0.12 µM $N_{14}$, 0.06 µM $N_{15}$, and 0.03 µM $N_{16}$), 200 mM 1-methylimidazole (1MI), 400 mM 2-methylbutyraldehyde (2MBA), 400 mM methyl isocyanide (MeNC), and 50 mM $MgCl_2$ incubated for 48 hours at 23°C | The oligo mix was prepared in advance as a 5 mM stock solution. Ligation-only conditions with 1-methylimidazole as the activating group | S10, 4 |
| **EEA_18NL(2, 3, ..., or 8) & EEB_18NL(9, 10, ..., or 16)** | Sequencing hairpin template and initially unactivated random-sequence oligonucleotides of increasing lengths ($pN_{2-16}$) in decreasing concentrations exposed to 23°C high concentration MeNC activation for 48 hours with 200 mM 1MI | 100 mM HEPES-Na pH 8, 1 µM 18N sequencing hairpin, 1.2 µM 5′ Handle Block oligo, 5′-phosphorylated random-sequence oligonucleotide, one length oligonucleotide at a time (500 µM $pN_2$; 250 µM $pN_3$; 125 µM $pN_4$; 62.5 µM $pN_5$; 31.3 µM $pN_6$; 15.6 µM $pN_7$; 7.8 µM $pN_8$; 3.9 µM $pN_9$; 2 µM $pN_{10}$; 0.98 µM $pN_{11}$; 0.49 µM $pN_{12}$; 0.24 µM $pN_{13}$; 0.12 µM $pN_{14}$; 0.06 µM $pN_{15}$; or 0.03 µM $pN_{16}$), 200 mM 1-methylimidazole (1MI), 400 mM 2-methylbutyraldehyde (2MBA), 400 mM methyl isocyanide (MeNC), and 50 mM $MgCl_2$ incubated for 48 hours at 23°C | Ligation-only conditions with 1-methylimidazole as the activating group | ST1, S9, S10, 4, S11 |
| **FF_18N1a** | Sequencing hairpin template, mixture of initially unactivated random-sequence oligonucleotides of increasing | 50 mM HEPES-Na pH 8, 1 µM 18N sequencing hairpin, 1.2 µM 5′ Handle Block oligo, 1 mM total oligonucleotide mixture (oligo mix) (500 µM $N_2$, 250 µM $N_3$, 125 µM | The oligo mix was prepared in advance as a 5 mM stock solution. | S11A |

| | | | | |
|---|---|---|---|---|
| | lengths ($pN_{2-16}$) in decreasing concentrations, and initially unactivated nucleotides exposed to four rounds of eutectic MeNC activation at 12 hour intervals followed by six rounds of 23°C MeNC activation at 2 hour intervals | $N_4$, 62.5 µM $N_5$, 31.3 µM $N_6$, 15.6 µM $N_7$, 7.81 µM $N_8$, 3.91 µM $N_9$, 1.95 µM $N_{10}$, 0.98 µM $N_{11}$, 0.49 µM $N_{12}$, 0.24 µM $N_{13}$, 0.12 µM $N_{14}$, 0.06 µM $N_{15}$, and 0.03 µM $N_{16}$), 20 mM rNMP (5 mM each of rAMP, rUMP, rGMP, and rCMP), 20.5 mM 2-aminoimidazole (2AI), and 30 mM $MgCl_2$; 200 mM total 2-methylbutyraldehyde (2MBA) added once + 50 mM methyl isocyanide (MeNC) added at each brief thaw between four frozen incubations (4 x 12 hours at -17°C); thaw + (200 mM total 2MBA added once + 100 mM MeNC added twice (2 x 2 hours at 23°C)) x 3; thaw + 18 hours at 23°C | | |
| **FF_18N1aflt** | Sequencing hairpin template, mixture of initially unactivated random-sequence oligonucleotides of increasing lengths ($pN_{2-16}$) at equal concentrations, and initially unactivated nucleotides exposed to four rounds of eutectic MeNC activation at 12 hour intervals followed by six rounds of 23°C MeNC activation at 2 hour intervals | 50 mM HEPES-Na pH 8, 1 µM 18N sequencing hairpin, 1.2 µM 5′ Handle Block oligo, 16 µM total oligonucleotide mixture (1 µM each of $pN_{2-16}$), 20 mM rNMP (5 mM each of rAMP, rUMP, rGMP, and rCMP), 20.5 mM 2-aminoimidazole (2AI), and 30 mM $MgCl_2$; 200 mM total 2-methylbutyraldehyde (2MBA) added once + 50 mM methyl isocyanide (MeNC) added at each brief thaw between four frozen incubations (4 x 12 hours at -17°C); thaw + (200 mM total 2MBA added once + 100 mM MeNC added twice (2 x 2 hours at 23°C)) x 3; thaw + 18 hours at 23°C | The oligonucleotide mixture was prepared in advance as a 100 µM stock solution. | S11B |
| **GG_18N26** | Sequencing hairpin template, mixture of initially unactivated random-sequence oligonucleotides of lengths 2-6 in decreasing concentrations, and initially unactivated nucleotides exposed to four rounds of eutectic MeNC activation followed by four rounds of 23°C MeNC activation at 24 hour intervals | 50 mM HEPES-Na pH 8, 1 µM 18N sequencing hairpin, 1.2 µM 5′ Handle Block oligo, 969 µM total oligonucleotide mixture (500 µM $N_2$, 250 µM $N_3$, 125 µM $N_4$, 62.5 µM $N_5$, 31.3 µM $N_6$), 20 mM rNMP (5 mM each of rAMP, rUMP, rGMP, and rCMP), 20.5 mM 2-aminoimidazole (2AI), and 30 mM $MgCl_2$; 200 mM total 2-methylbutyraldehyde (2MBA) added once + 50 mM methyl isocyanide (MeNC) added at each brief thaw between | The oligonucleotide mixture was prepared in advance as a 5x stock solution. | S11C |

| | | | | |
|---|---|---|---|---|
| | | four frozen incubations (4 x 24 hours at -17°C); thaw + 24 hours at 23°C; 200 mM total 2MBA added once + 50 mM MeNC added four times (4 x 24 hours at 23°C) | | |
| **GG_18N716** | Sequencing hairpin template, mixture of initially unactivated random-sequence oligonucleotides of lengths 7-16 in decreasing concentrations, and initially unactivated nucleotides exposed to four rounds of eutectic MeNC activation followed by four rounds of 23°C MeNC activation at 24 hour intervals | 50 mM HEPES-Na pH 8, 1 μM 18N sequencing hairpin, 1.2 μM 5′ Handle Block oligo, 31 μM total oligonucleotide mixture (15.6 μM $N_7$, 7.81 μM $N_8$, 3.91 μM $N_9$, 1.95 μM $N_{10}$, 0.98 μM $N_{11}$, 0.49 μM $N_{12}$, 0.24 μM $N_{13}$, 0.12 μM $N_{14}$, 0.06 μM $N_{15}$, and 0.03 μM $N_{16}$), 20 mM rNMP (5 mM each of rAMP, rUMP, rGMP, and rCMP), 20.5 mM 2-aminoimidazole (2AI), and 30 mM $MgCl_2$; 200 mM total 2-methylbutyraldehyde (2MBA) added once + 50 mM methyl isocyanide (MeNC) added at each brief thaw between four frozen incubations (4 x 24 hours at -17°C); thaw + 24 hours at 23°C; 200 mM total 2MBA added once + 50 mM MeNC added four times (4 x 24 hours at 23°C) | The oligonucleotide mixture was prepared in advance as a 5x stock solution. | S11D |
| **GG_18N4deg** | Sequencing hairpin template, mixture of initially unactivated random-sequence oligonucleotides of lengths 2-6 in decreasing concentrations, and initially unactivated nucleotides exposed to four rounds of eutectic MeNC activation followed by four rounds of 4°C MeNC activation at 24 hour intervals | 50 mM HEPES-Na pH 8, 1 μM 18N sequencing hairpin, 1.2 μM 5′ Handle Block oligo, 969 μM total oligonucleotide mixture (500 μM $N_2$, 250 μM $N_3$, 125 μM $N_4$, 62.5 μM $N_5$, 31.3 μM $N_6$), 20 mM rNMP (5 mM each of rAMP, rUMP, rGMP, and rCMP), 20.5 mM 2-aminoimidazole (2AI), and 30 mM $MgCl_2$; 200 mM total 2-methylbutyraldehyde (2MBA) added once + 50 mM methyl isocyanide (MeNC) added at each brief thaw between four frozen incubations (4 x 24 hours at -17°C); thaw + 24 hours at 4°C; 200 mM total 2MBA added once + 50 mM MeNC added four times (4 x 24 hours at 4°C) | The oligonucleotide mixture was prepared in advance as a 5x stock solution. | S11E |
| **GG_18N264** | Sequencing hairpin template, mixture of initially unactivated random-sequence oligonucleotides of lengths 2-6 in | 50 mM HEPES-Na pH 8, 1 μM 18N sequencing hairpin, 1.2 μM 5′ Handle Block oligo, 969 μM total oligonucleotide mixture (500 μM $N_2$, 250 μM $N_3$, 125 μM $N_4$, 62.5 μM | The oligonucleotide mixture was prepared in advance as a 5x stock solution. | S11F |

| | | | | |
|---|---|---|---|---|
| | decreasing concentrations, and initially unactivated nucleotides exposed to four rounds of eutectic MeNC activation at 12 hour intervals followed by four rounds of 23°C MeNC activation at 24 hour intervals | $N_5$, 31.3 µM $N_6$), 20 mM rNMP (5 mM each of rAMP, rUMP, rGMP, and rCMP), 20.5 mM 2-aminoimidazole (2AI), and 30 mM $MgCl_2$; 200 mM total 2-methylbutyraldehyde (2MBA) added once + 50 mM methyl isocyanide (MeNC) added at each brief thaw between four frozen incubations (4 x 12 hours at -17°C); thaw + 24 hours at 23°C; 200 mM total 2MBA added once + 50 mM MeNC added four times (4 x 24 hours at 23°C) | | |
| **GG_18N265** | Sequencing hairpin template, mixture of initially unactivated random-sequence oligonucleotides of lengths 2-6 in decreasing concentrations, and initially unactivated nucleotides exposed to four rounds of eutectic MeNC activation at 12 hour intervals followed by four rounds of 4°C MeNC activation at 24 hour intervals | 50 mM HEPES-Na pH 8, 1 µM 18N sequencing hairpin, 1.2 µM 5′ Handle Block oligo, 969 µM total oligonucleotide mixture (500 µM $N_2$, 250 µM $N_3$, 125 µM $N_4$, 62.5 µM $N_5$, 31.3 µM $N_6$), 20 mM rNMP (5 mM each of rAMP, rUMP, rGMP, and rCMP), 20.5 mM 2-aminoimidazole (2AI), and 30 mM $MgCl_2$; 200 mM total 2-methylbutyraldehyde (2MBA) added once + 50 mM methyl isocyanide (MeNC) added at each brief thaw between four frozen incubations (4 x 12 hours at -17°C); thaw + 24 hours at 4°C; 200 mM total 2MBA added once + 50 mM MeNC added four times (4 x 24 hours at 4°C) | The oligonucleotide mixture was prepared in advance as a 5x stock solution. | S11G |
| **GG_18N266** | Sequencing hairpin template, mixture of initially unactivated random-sequence oligonucleotides of lengths 2-6 in decreasing concentrations, and initially unactivated nucleotides exposed to four rounds of eutectic MeNC activation at 12 hour intervals followed by four rounds of 23°C MeNC activation at 12 hour intervals | 50 mM HEPES-Na pH 8, 1 µM 18N sequencing hairpin, 1.2 µM 5′ Handle Block oligo, 969 µM total oligonucleotide mixture (500 µM $N_2$, 250 µM $N_3$, 125 µM $N_4$, 62.5 µM $N_5$, 31.3 µM $N_6$), 20 mM rNMP (5 mM each of rAMP, rUMP, rGMP, and rCMP), 20.5 mM 2-aminoimidazole (2AI), and 30 mM $MgCl_2$; 200 mM total 2-methylbutyraldehyde (2MBA) added once + 50 mM methyl isocyanide (MeNC) added at each brief thaw between four frozen incubations (4 x 12 hours at -17°C); thaw + 24 hours at 23°C; 200 mM total 2MBA added once + 50 mM MeNC added four times (4 x 12 hours at 23°C) | The oligonucleotide mixture was prepared in advance as a 5x stock solution. | S11H |

| | | | | |
|---|---|---|---|---|
| **GG_18N267** | Sequencing hairpin template, mixture of initially unactivated random-sequence oligonucleotides of lengths 2-6 in decreasing concentrations, and initially unactivated nucleotides exposed to four rounds of eutectic MeNC activation at 12 hour intervals followed by four rounds of 4°C MeNC activation at 12 hour intervals | 50 mM HEPES-Na pH 8, 1 µM 18N sequencing hairpin, 1.2 µM 5′ Handle Block oligo, 969 µM total oligonucleotide mixture (500 µM $N_2$, 250 µM $N_3$, 125 µM $N_4$, 62.5 µM $N_5$, 31.3 µM $N_6$), 20 mM rNMP (5 mM each of rAMP, rUMP, rGMP, and rCMP), 20.5 mM 2-aminoimidazole (2AI), and 30 mM $MgCl_2$; 200 mM total 2-methylbutyraldehyde (2MBA) added once + 50 mM methyl isocyanide (MeNC) added at each brief thaw between four frozen incubations (4 x 12 hours at -17°C); thaw + 24 hours at 4°C; 200 mM total 2MBA added once + 50 mM MeNC added four times (4 x 12 hours at 4°C) | The oligonucleotide mixture was prepared in advance as a 5x stock solution. | S11I |
| **HH_18N4r** | (Sequencing hairpin template, mixture of initially unactivated random-sequence oligonucleotides of increasing lengths ($pN_{2-16}$) in decreasing concentrations, and initially unactivated nucleotides exposed to four rounds of eutectic MeNC activation followed by four rounds of 23°C MeNC activation at 24 hour intervals) x 4 | [50 mM HEPES-Na pH 8, 1.5 µM 18N sequencing hairpin, 1.8 µM 5′ Handle Block oligo, 1 mM total oligonucleotide mixture (oligo mix) (500 µM $N_2$, 250 µM $N_3$, 125 µM $N_4$, 62.5 µM $N_5$, 31.3 µM $N_6$, 15.6 µM $N_7$, 7.81 µM $N_8$, 3.91 µM $N_9$, 1.95 µM $N_{10}$, 0.98 µM $N_{11}$, 0.49 µM $N_{12}$, 0.24 µM $N_{13}$, 0.12 µM $N_{14}$, 0.06 µM $N_{15}$, and 0.03 µM $N_{16}$), 20 mM rNMP (5 mM each of rAMP, rUMP, rGMP, and rCMP), 20.5 mM 2-aminoimidazole (2AI), and 30 mM $MgCl_2$; 260 mM total 2-methylbutyraldehyde (2MBA) added once + 50 mM methyl isocyanide (MeNC) added at each brief thaw between four frozen incubations (4 x 24 hours at -17°C); thaw + 24 hours at 23°C; 260 mM total 2MBA added once + 50 mM MeNC added four times (4 x 24 hours at 23°C)]; quench, 18N sequencing hairpin and 5′ Handle Block oligo purified, and entire reaction repeated (4 rounds total) | This experiment is equivalent to CCN_18N1a, repeated four times, except a higher initial concentration of the sequencing hairpin and the 5′ Handle Block were used to account for substrate loss at each quench / purification step. The experiment took 40 days. The oligo mix was prepared in advance as a 5 mM stock solution. | S13 |
| **HH_18N6r** | (Sequencing hairpin template, mixture of initially unactivated random-sequence oligonucleotides of increasing | [50 mM HEPES-Na pH 8, 2 µM 18N sequencing hairpin, 2.4 µM 5′ Handle Block oligo, 1 mM total oligonucleotide mixture (oligo mix) (500 µM $N_2$, 250 µM $N_3$, 125 µM | This experiment is equivalent to CCN_18N1a, repeated six times, except a | 5, S13, S14 |

| | | | |
|---|---|---|---|
| | lengths ($pN_{2-16}$) in decreasing concentrations, and initially unactivated nucleotides exposed to four rounds of eutectic MeNC activation followed by four rounds of 23°C MeNC activation at 24 hour intervals) x 6 | $N_4$, 62.5 µM $N_5$, 31.3 µM $N_6$, 15.6 µM $N_7$, 7.81 µM $N_8$, 3.91 µM $N_9$, 1.95 µM $N_{10}$, 0.98 µM $N_{11}$, 0.49 µM $N_{12}$, 0.24 µM $N_{13}$, 0.12 µM $N_{14}$, 0.06 µM $N_{15}$, and 0.03 µM $N_{16}$), 20 mM rNMP (5 mM each of rAMP, rUMP, rGMP, and rCMP), 20.5 mM 2-aminoimidazole (2AI), and 30 mM $MgCl_2$; 260 mM total 2-methylbutyraldehyde (2MBA) added once + 50 mM methyl isocyanide (MeNC) added at each brief thaw between four frozen incubations (4 x 24 hours at -17°C); thaw + 24 hours at 23°C; 260 mM total 2MBA added once + 50 mM MeNC added four times (4 x 24 hours at 23°C)]; quench, 18N sequencing hairpin and 5′ Handle Block oligo purified, and entire reaction repeated (6 rounds total) | higher initial concentration of the sequencing hairpin and the 5′ Handle Block were used to account for substrate loss at each quench / purification step. The experiment took 60 days. The oligo mix was prepared in advance as a 5 mM stock solution. |
